# Novel childhood experience suggests eccentricity drives organization of human visual cortex

**DOI:** 10.1101/415729

**Authors:** Jesse Gomez, Michael Barnett, Kalanit Grill-Spector

## Abstract

The functional organization of human high-level visual cortex, such as face and place-selective regions, is strikingly consistent across individuals. A fundamental, unanswered question in neuroscience is what dimensions of visual information constrain the development and topography of this shared brain organization? To answer this question, we scanned with fMRI a unique group of adults who, as children, engaged in extensive experience with a novel stimulus–Pokémon–which are dissimilar from other ecological categories such as faces and places along critical dimensions (foveal bias, rectilinearity, size, animacy) from. We find that experienced adults not only demonstrate distinct and consistent distributed cortical responses to Pokémon, but their activations suggest that it is the experienced retinal eccentricity during childhood that predicts the locus of distributed responses to Pokémon in adulthood. These data advance our understanding about how childhood experience and functional constraints shape the functional organization of the human brain.

## Introduction

Humans possess the remarkable ability to rapidly recognize a wide array of visual stimuli. This ability is thought to occur from cortical computations in the ventral visual stream^1^: a processing hierarchy extending from primary visual cortex (V1) to ventral temporal cortex (VTC). Prior research has shown that VTC responses are key for visual recognition because (i) distributed VTC responses contain information about objects^2–4^ and categories^5^, and (ii) responses in category-selective regions within VTC such as face-, body-, word-, and place-selective regions^6–12^, are linked to perception of these categories^13–15^. Notably, distributed VTC response patterns to different visual categories are distinct from one another and are arranged with remarkable spatial consistency along the cortical sheet across individuals^16–23^. For example, peaks in distributed VTC response patterns to faces are consistently found on the lateral fusiform gyrus (FG). While several theories have been suggested for the consistent spatial topography of VTC^24–26^, developmental studies suggest that experience may be key for normal development of VTC and recognition abilities. For example, behavioral studies suggest that typical development of recognition abilities is thought to be reliant on viewing experience during childhood^27–32^. However, the nature of childhood experience that leads to the consistent spatial functional topography of VTC, whether it is the way stimuli such as faces or places are viewed, or the image-level statistics of the stimuli themselves, remains unknown.

Several theories have proposed attributes that may underlie the functional topography of human high-level visual cortex. These include: (1) Eccentricity bias of retinal images associated with typical viewing of specific categories^19,24^; e.g., face discrimination is thought to require high visual acuity supported by foveal vision, and places are thought to be processed more with peripheral vision as in the real world they occupy the entirety of the visual field. (2) Average rectilinearity of stimuli of particular categories ^25,33^; e.g., faces are curvilinear, but manmade places tend to be rectilinear. (3) The perceived animacy of stimuli^5,34–37^; e.g., faces are perceived to be animate whereas places are not, and (4) real-world size of stimuli^38^; e.g., faces are physically smaller than places and buildings.

These theories each propose an underlying principle describing the coarse functional topography of VTC relative to its cortical macroanatomy. That is, inherent in all these theories is the idea that a physical or perceived dimension of a stimulus maps to a linear dimension along the cortical surface. For example, in human VTC, small, curvy, animate, and foveal stimuli elicit stronger responses lateral to the mid-fusiform sulcus (MFS), while large, linear, inanimate, and peripherally-extending stimuli elicit stronger responses in cortex medial to the MFS. However, which of these dimensions drives the development of the functional organization of VTC is unknown.

Research on cortical plasticity in animals has made two key discoveries related to this question. First, of the aforementioned attributes, eccentricity representations in early and intermediate visual cortex are likely established in infancy^39,40^, as they may be constrained by both wiring^41^ and neural activity that starts *in utero*^42^. For example, research of ferret and mouse development suggests that retinal waves during gestation and before eye-opening are sufficient to establish eccentricity representations in visual cortex^40^. Additionally, an eccentricity proto-architecture is detectable early in macaque development^39^. Second, visual development has a critical period during which the brain is particularly malleable and sensitive to visual experience^32,43–48^. For example, prior research in macaques suggests that new category representations in high-level visual cortex emerge with experience only early in development, but not if the same experience happens in adulthood^43^. Further, visual deprivation to a category (e.g., faces) in infancy results in a lack of development of a cortical representation for that category^32^. Together these findings make the following predictions regarding human development: First, if eccentricity representations in high-level visual cortex are present early in development, then eccentricity stands to be a strong developmental constraint for the later emergence of object representations. Second, testing theories of VTC development requires measuring the effects of childhood experience on the formation of new brain representations.

Thus, to investigate the developmental origins of the functional organization of VTC requires testing the effects of exhaustive childhood experience with a novel visual category that is distinct along the four aforementioned visual dimensions from natural categories. Performing a sufficient experimental manipulation in a laboratory setting with children would require an unfeasible length of time. However, there exist adults who, in their childhood, had prolonged, rewarded, and crucially similar visual experience with a shared novel stimulus category. In 1996, Nintendo released the popular game, Pokémon, alongside a handheld playing device, the GameBoy. Children as young as five years old played this game, in which individuating the animal-like creatures of Pokémon was integral to optimal game performance. Furthermore, game playing conditions were nearly identical across individuals: children held the device at a similar, arm’s length, viewing distance, repeatedly for hours a day, over the period of years. Importantly, Pokémon were rendered in a small 2.5 × 2.5 cm region of the GameBoy screen with large pixels, making Pokémon, compared to faces or places, a unique visual category that is of a constant size and viewed with foveal vision, but possesses strong linear features. While many Pokémon characters resemble animals, and children saw Pokémon animations, they were never encountered in the real world. Thus, Pokémon’s animacy and real-world size are inferred attributes rather than physical ones. As Pokémon are unique from both faces and places along these visual dimensions, we can use them to answer a fundamental question: Does prolonged experience individuating Pokémon result in novel information representation in VTC, and moreover, is the location of this response predicted by a specific visual dimension?

To answer these questions, we scanned with functional magnetic resonance imaging (fMRI) 11 experienced adult participants who began playing Pokémon between the ages of 5 and 8 years old, in addition to 11 age-matched adult novices inexperienced with Pokémon. During fMRI participants viewed stimuli from eight categories: faces, animals, cartoons, bodies, words, cars, corridors, and Pokémon (**Fig 1A**). We performed multi-voxel pattern analysis (MVPA) of VTC responses to these stimuli in each subject, combined with decoding analyses, to assess (i) if prolonged experience results in an informative representation for Pokémon in experienced versus novice participants, and (ii) if distributed VTC representations form a consistent spatial topography across VTC in experienced versus novice participants. Should experienced individuals demonstrate an informative and spatially consistent representation for Pokémon compared to other categories, it would afford us the opportunity to ask if the topography of this representation across VTC is predicted by one of the four visual dimensions (eccentricity bias, rectilinearity, animacy, and perceived size), which we quantified for Pokémon, faces, and places.

**Figure 1:**
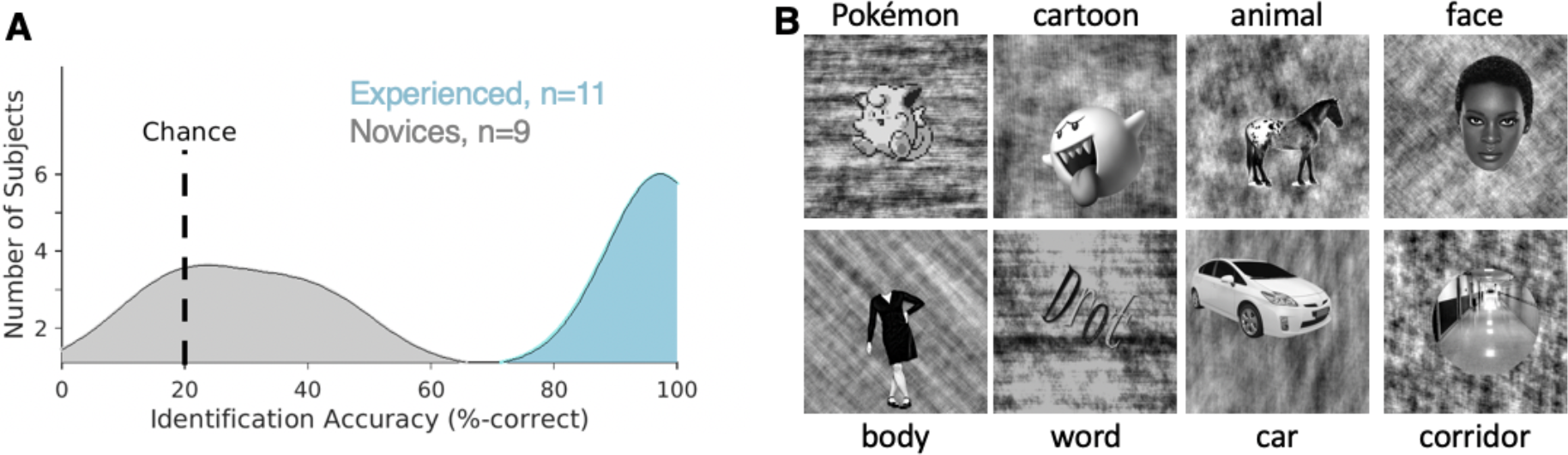
Localizer stimuli and behavioral naming performance. **A)** Histograms of subject accuracies (% correct) from a 5-alternative-choice Pokémon naming task outside the scanner. Experienced subjects (blue) significantly outperformed novices (gray). **B)** Example stimuli from each of the categories used in the fMRI experiment. In each 4 s trial, subjects viewed 8 different stimuli from each category a rate of 2 Hz while performing an Oddball task to detect a phase-scrambled stimulus With no intact Object overlaid. Subjects participated 6 runs, 3 mins and 38 s each, using different stimuli.

## Results

### Childhood experience with Pokémon results in distinct and reproducible information across VTC

Experienced participants were adults (n=11, mean age 24.3±32.8 years, 3 female) initially chosen through self-report, who began playing Pokémon between the ages of 5 and 8. Experienced participants were included in the study if they continued to play the game throughout childhood, and revisited playing the game as adults. Novice participants were chosen as similarly-aged and educated adults who never played Pokémon (n=11, mean age 29.5±5.4, 7 female). To validate their self-report, participants completed a behavioral experiment in which they viewed 40 Pokémon images from the original Nintendo game, and for each image identified its name from 5-multiple-choices. Experienced participants significantly outperformed novices in naming ability of Pokémon (t(18)=18.2, p<10^−10^; **Fig 1A**). Despite not being able to name Pokémon, novices are capable of visually distinguishing and individuating Pokémon characters (**Fig S1**)

All participants underwent fMRI while viewing faces, bodies, cartoons, pseudowords, Pokémon, animals, cars, and corridors (**Fig 1B**). Cartoons and animals were chosen to create a strict comparison to the Pokémon stimuli, and other categories were included as they have well established and reproducible spatial topography across VTC. Stimuli were randomly presented at a rate of 2 Hz, in 4 s blocks, each containing 8 images from a category. Participants performed an oddball detection task to ensure continuous attention throughout the scan. Subjects participated in 6 runs with different stimuli from these categories.

We first examined if childhood experience affects the representation of category information in VTC, which was anatomically defined in each subject’s native brain (Methods). Thus, we measured in each participant the representational similarity among distributed VTC responses to the 8 categories across runs. Each cell in the representational similarity matrix (RSM) is the voxelwise correlation between the distributed VTC responses to different images of the same category (diagonal) or different categories (off-diagonal) across split halves of the data. Then, we averaged the RSMs across participants of each group and compared across groups.

We hypothesized that representation similarity of distributed VTC responses will have one of four outcomes. 1) *Null hypothesis:* Pokémon will not elicit a consistent response pattern in VTC in any group and will have near-zero correlation with itself and other categories. 2) *Animate hypothesis:* Pokémon, which have faces, limbs, and resemble animals to some extent, will have positive correlations with animate categories such as faces, bodies, and animals. 3) *Expertise hypothesis:* If Pokémon are processed as a category of expertise, then distributed responses to Pokémon will be most correlated with distributed responses to faces, as the expertise hypothesis predicts that expert stimuli are processed in face-selective regions^49^. 4) *Distinctiveness hypothesis*: as Pokémon constitute a category of their own, they will elicit a response pattern that is unique from other categories. Thus, correlations among distributed responses to different Pokémon will be positive and substantially higher than the correlation between Pokémon and items of other categories.

Illustrated in the RSM in **Fig 2A**, experienced participants differ markedly from novices in their distributed VTC response patterns to Pokémon. Unlike novices, who demonstrate little to no reproducible pattern for Pokémon in VTC consistent with the null hypothesis (mean correlation±std, r=0.1±0.06), experienced participants instead demonstrate a significantly more reproducible response pattern for Pokémon (r=0.27±0.11; significant between-group difference: t(20)=4, p<0.0008). Additionally, distributed responses to Pokémon were distinct from those of other categories in experienced participants. We quantify this effect by calculating the mean dissimilarity (D=1-r) of distributed responses to Pokémon from other categories. Distributed responses to Pokémon are significantly more dissimilar from distributed responses to the other categories in experienced participants than controls (t(20)=4.4, p<0.0005). Interestingly, in experienced participants, Pokémon response patterns are significantly dissimilar (*ts*(20)<4.2, *ps*<0.0005) from those of faces (D±std: 0.97±0.08), bodies (1.1±0.08), and animals (0.9±0.07) despite Pokémon having faces, bodies, and animal-like features themselves. In contrast, when excluding Pokémon, groups are not significantly different in the mean dissimilarity between distributed responses to other pairs of categories (t(20)=0.52, p=0.6).

**Figure 2:**
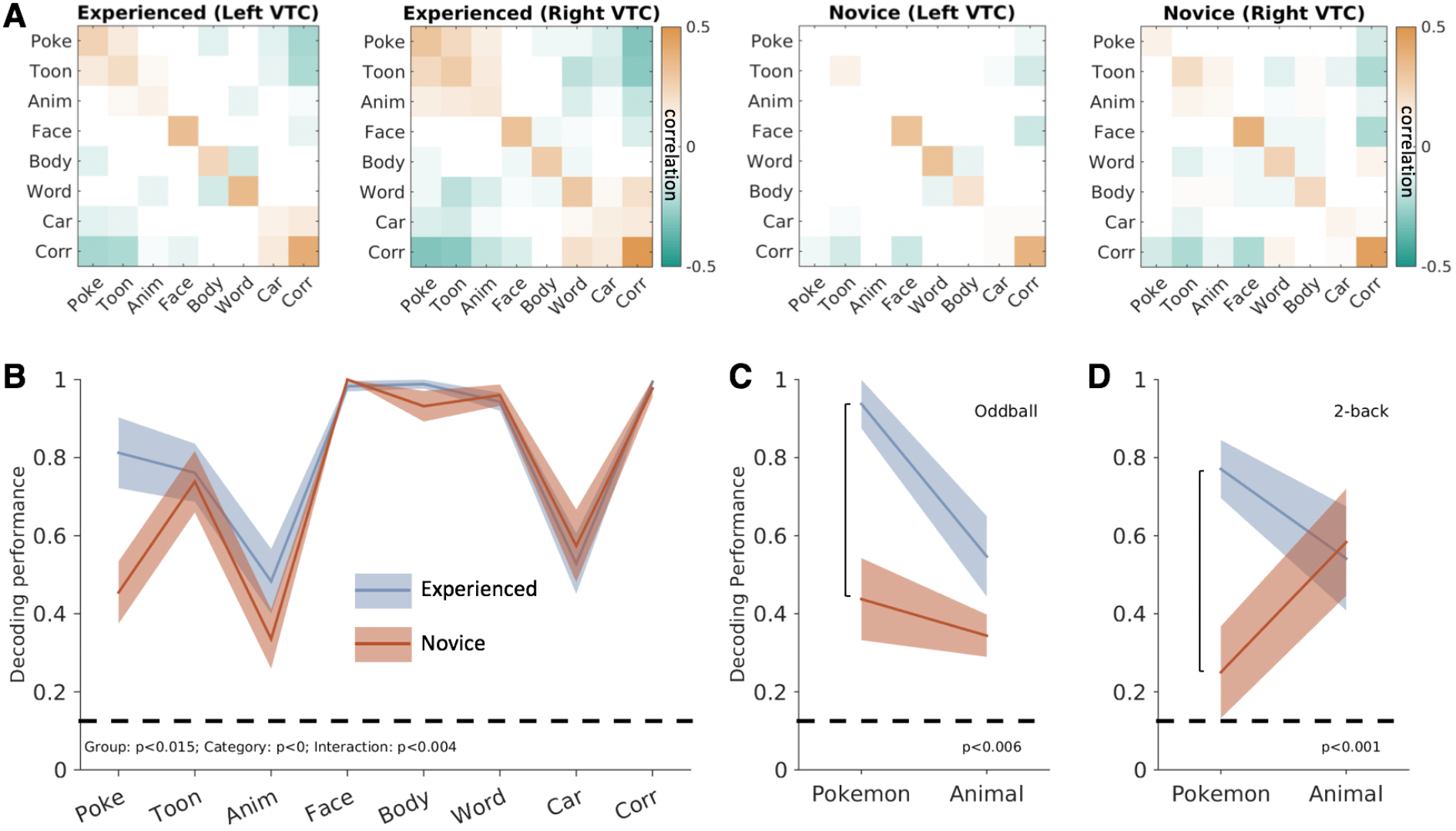
Experienced participants demonstrate consistent and distinct representation for Pokémon compared to novices. **A)** Representational Similarity matrices (RSMS) calculated by correlating distributed responses (z-scored voxel betas) from an anatomical VTC ROI split-halves of the fMRI experiment. Positive values are presented in orange, negative values in green, and near-zero values are white. **B)** Decoding performance from the winner-take-all classifier trained and tested on split-halves of the fMRI data from bilateral VTC. Experienced subjects (n=11) are in blue, novices (n=11) are in orange. *Shaded region:* standard error across subjects within a group. *Dashed line:* chance level performance. Decoding performance is represented as a fraction of 1, with 1 corresponding to 100% decoding accuracy P-values indicate significant grouping factors from an ANOVA. **(C-D)** Decoding performance from distributed bilateral VTC responses for experienced (n=4) and novice (n=5) subjects who were brought back to undergo an additional fMRI experiment with an 2-back task. Same subjects are shown in C and D. P-values indicate significant t-test comparing group decoding differences for Pokémon. **C)** Decoding performance from the original oddball task. **D)** Decoding performance during the 2-back task.

To quantify the information content in VTC and test if it varies with experience, we constructed a winner-take-all classifier trained and tested on independent halves of the data to determine if the category of the stimulus can be classified based on distributed VTC response patterns, and if this performance depends on childhood experience. While classification in both groups was above chance for all categories, performance varied by category and group (**Fig 2B**). An analysis of variance (ANOVA) with factors of stimulus category and participant group (experienced/novices) on classification performance revealed significant effects of stimulus category (F(7,160)=36.5, *p*<0.0001), group (F(1,160)=6, *p*=0.015), and a significant interaction between group and category (F(7,160)=3.1, *p*<0.005). Importantly, this interaction was driven by differential classification of brain responses to Pokémon across groups: in experienced participants performance was 81.3%±9% (mean±SE), which was significantly higher (post hoc t-test, p<0.002), and almost double that of the 45%±8% classification in novices (**Fig 2B**). Classification performance was also numerically higher for bodies and animals in experienced compared to novice participants, but this difference, along with classification performance for other stimuli, were not significantly different between groups (**Fig 2C**). Lastly, while there were more male experienced participants and more female novices, behavioral naming of Pokémon as well as decoding accuracy was not higher in experienced males than in experienced females (**Fig S2B,C**). These analyses suggest that childhood experience with Pokémon generates a reliable and informative distributed representation of Pokémon in VTC.

Can differences in attention account for this pattern of results? While it has been argued that attention can boost signals to the category of expertise^50^, others have argued that attention to the expert category does not explain the enhanced cortical activity to viewing it^51^. Furthermore, it is unclear if attention alone could induce a distinct distributed response that could be reliably decoded. Nonetheless, to test the possibility that selective attention to Pokémon in experienced participants is driving the improved classification performance, we invited a subset of experienced and novice participants to perform an additional fMRI experiment with the same stimuli, using a different task that was attention-demanding and equally so across all categories. Here, participants performed a 2-back task, indicating whether an image repeated after an intervening image. In this second fMRI experiment, classifying Pokémon from distributed VTC responses was again significantly higher in experienced (77%±14%) compared to novice participants (25%±23%, between-group difference: t(7)=5.6, *p*<0.001, **Fig 2D**). In contrast, controlling attention generated similar classification of animals across groups (**Fig 2D**). These data suggest that differential attention across groups to Pokémon is not the driving factor leading to the distinct and reproducible representation of Pokémon stimuli in the VTC of experienced participants.

### Visual features make different predictions for the emergent location of cortical responses

Results of the prior multivoxel pattern analyses suggest that intense childhood experience with a novel visual category results in a reproducible distributed response across human VTC that is distinct from other categories. An open question is if Pokémon generate distributed response patterns with similar topography across experienced participants. Thus, we generated statistical parametric maps contrasting the response to each category vs. all others (units of T-values), and compared across groups. Additionally, in typical adults, stimulus dimensions such as eccentricity^19,52^, animacy^37,53^, size^26^, and curvilinearity^25^ are mapped to a physical, lateral-medial axis across VTC^54^. As the topography of responses to faces on lateral VTC and places on medial VTC, generate the most differentiated topographies, we sought to analyze the properties of Pokémon stimuli for these attributes relative to faces and places. Thus, we use these metrics to generate predictions for what could be the emergent topography of a Pokémon representation in experienced participants.

As expected in both groups, unthresholded contrast maps demonstrating preference for faces and places showed the typical topography in relation to major anatomical landmarks. That is, despite both anatomical and functional variability between subjects, preference for faces was found in the lateral fusiform gyrus (FG) and preference for places in the collateral sulcus (CoS), as illustrated for example subjects in **Fig 3**. Striking differences, however, can be observed when examining VTC responses to Pokémon. In novices, Pokémon do not elicit preferential responses in VTC, like faces or corridors (**Fig 3A**). In contrast, an example experienced subject demonstrates robust preference to Pokémon in the lateral FG and occipitotemporal sulcus (OTS, **Fig 3B**). This pattern was readily observable in all other experienced participants (**Fig S3**). Given that these data suggest childhood experience with Pokémon result in a spatially consistent topography for Pokémon across individuals, we next asked: what attributes of Pokémon drive this topography?

**Figure 3:**
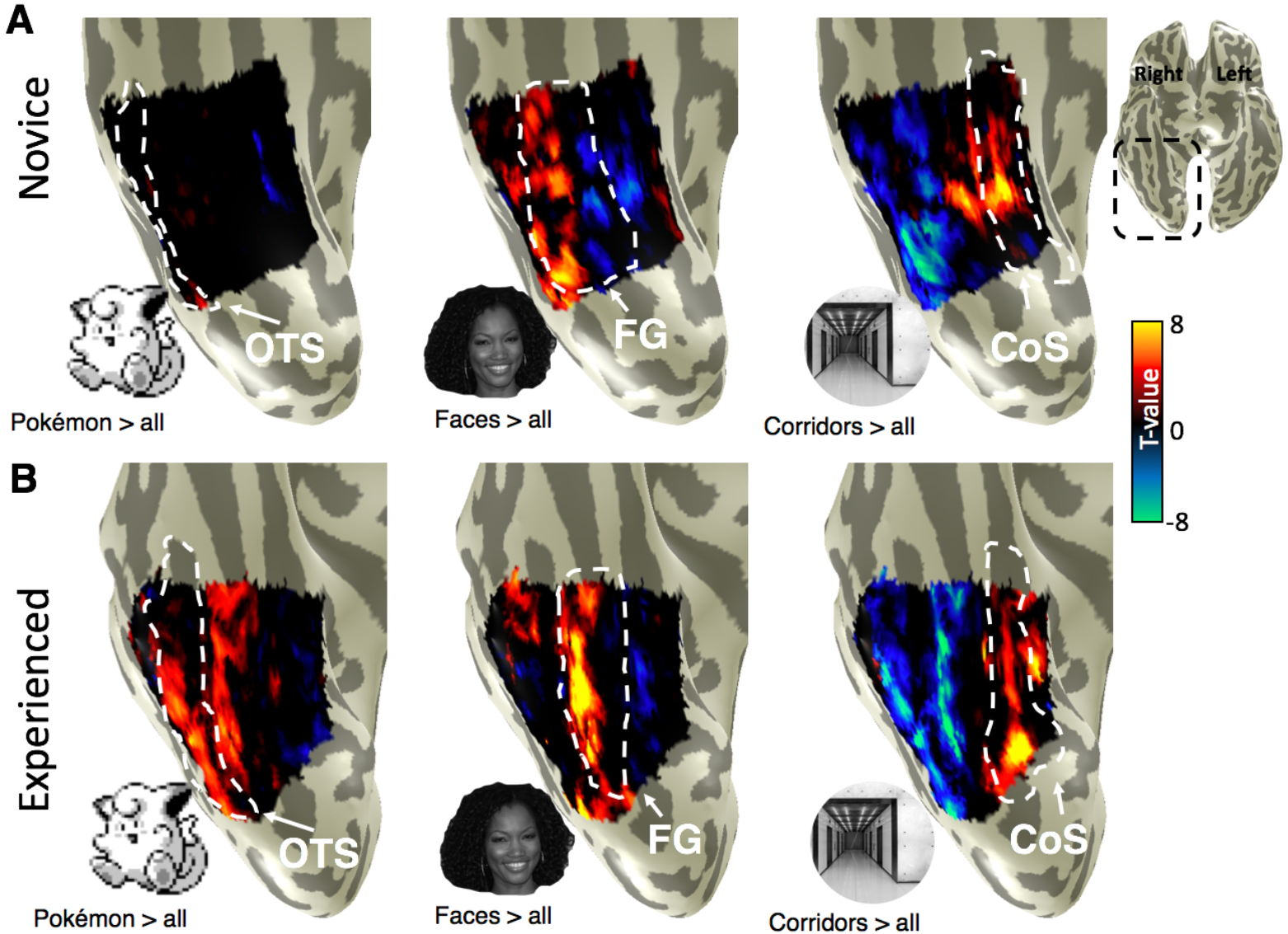
Distinct cortical representation for Pokémon in experienced subjects. **A)** Unthresholded parameter maps on the inflated ventral cortical surface zoomed on VTC (see inset) in an example novice subject (26yo female) for the contrasts of Pokémon (left), faces (center), and corridors (right), each versus all other stimuli. *White dotted outlines:* cortical folds of occipitotemporal sulcus (OTS, left), fusiform gyrus (FG, center), and collateral sulcus (CoS, right). **B)** Same as A, but for an example experienced subject (26yo male).

The stimuli of faces, corridors, and Pokémon used in the localizer experiments were submitted to a variety of analyses with the goal of ordering these categories linearly along different feature spaces (see Methods). Stimuli were analyzed for physical attributes of *foveal bias*, that is, the retinal size of images when fixated on across a range of typical viewing distances and *rectilinearity*, using the Rectilinearity Toolbox^25^, which evaluates the presence of linear and curved features at a range of spatial scales. Additionally, stimuli were rated for attributes of *size* and *animacy*, by independent raters.

We first evaluated image-based attributes. To estimate retinal size, we assumed that people foveate on the center of the item, and evaluated retinal image sizes as one would interact with these stimuli on a daily basis. In regards to retinal image size, Pokémon tend to span the central 2° when foveated upon on the Gameboy. The distribution of Pokémon retinal sizes partially overlapped, but were significantly smaller and thus more foveally biased, than faces (t(286)=15.7, p<10^−8^, **Fig 4A**). Both Pokémon and faces generate significantly smaller retinal images than a place category of corridors (*ts*>20, *p*s<10^−10^), which often occupy the entire visual field and thus generate large retinal images which extend to the peripheral visual field even when foveated upon. This metric predicts that if retinal image size ranking (from foveally biased to peripherally biased: Pokémon, faces, corridors) drives the generation of distributed responses, then Pokémon representations should fall in the most foveal representations, and places on the most peripheral ones. As the representation of retinal eccentricity in VTC moves from lateral (most foveal) to medial (most peripheral), this attribute predicts that emergent activations to Pokémon in experienced participants’ VTC should lie on the OTS, largely lateral to, but partially overlapping face-selective regions on the FG (**Fig 4A**).

**Figure 4:**
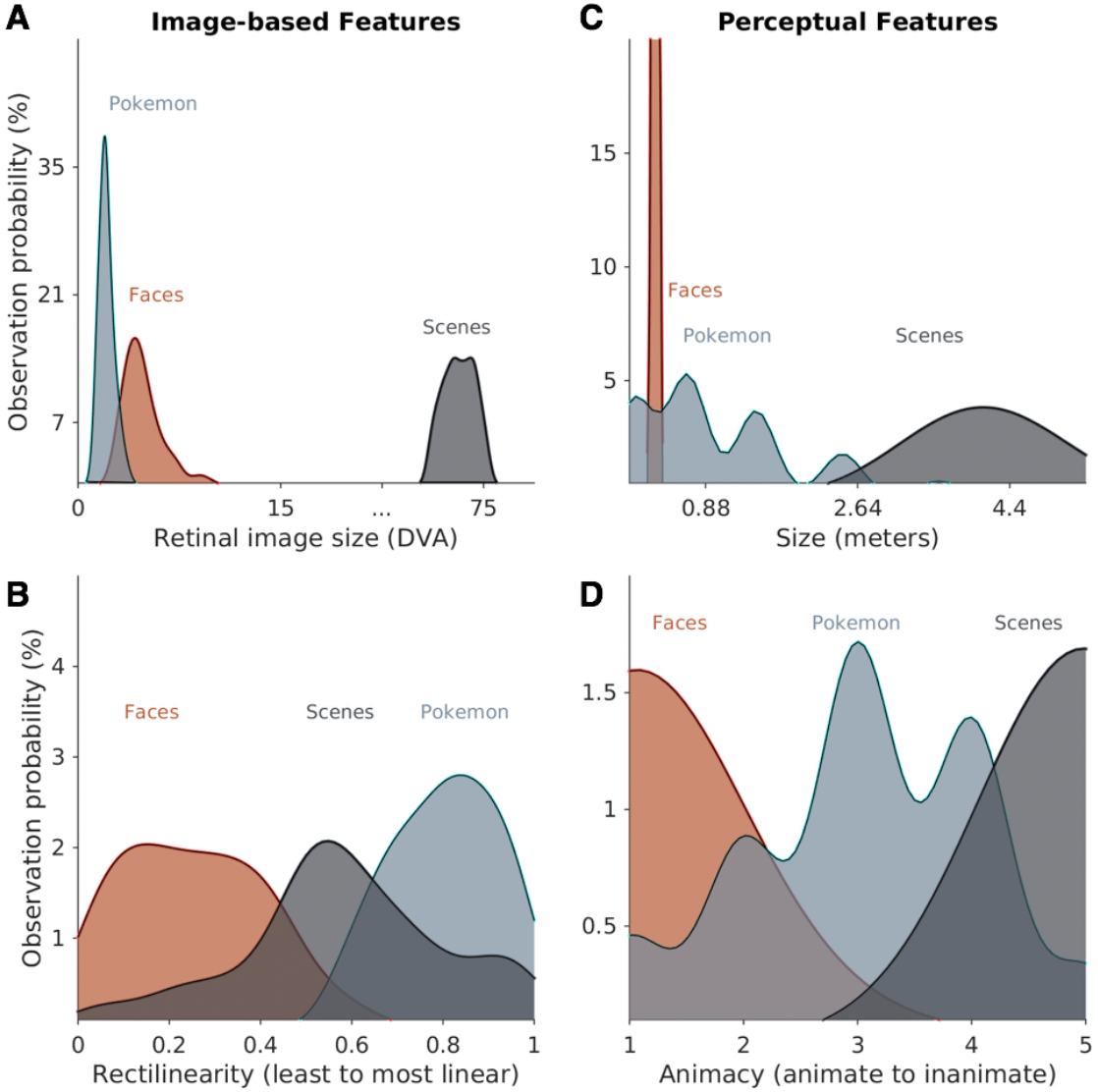
Different visual feature statistics predict different cortical locations for Pokémon. **A)** Histograms depicting the distributions of retinal image sizes produced by Pokémon (cyan), faces (orange), and corridors (gray) in a simulation varying viewing distance across a range of sample stimuli. *DVA:* degrees of visual angle. **B)** Histograms of the relative rectilinearity scores of faces, corridors, and Pokémon as measured using the wavelet filtering toolbox from Nasr, Echavarria, and Tootell (2014). **C)** Histograms depicting the distribution of the perceived physical size of Pokémon (from 28 raters), and distributions of the physical sizes of faces and corridor stimuli. The distributions of face and corridor size were produced using Gaussian distributions with standard deviations derived from either anatomical or physical variability within the stimulus category (see Methods). **D)** Histograms representing the scores of perceived animacy collected from a group of 42 independent raters who rated stimuli of faces, Pokémon, and corridors for how “living or animate” these stimuli were perceived to be.

Analyses of the rectilinearity of Pokémon, faces, and corridors shows that faces are the least linear of these 3 categories, and the original Pokémon stimuli, which were constructed from large square pixels, are the most rectilinear compared to both faces (t(383)=37.5,p<10^−15^) and corridors (t(298)=10.7, p<10^−10^). As the rectilinearity cortical axis is also arranged from curvy (lateral VTC) to rectilinear (medial VTC), rectilinearity predicts that preference to Pokémon in VTC should lie medial to both face- and place-selective cortex, potentially in the CoS or the parahippocampal gyrus (**Fig 4B**). We also evaluated *perceived* rectilinearity in a separate group of novices (**Fig S4**). While perceived rectilinearity situates Pokémon between faces (the most curvy) and corridors (perceived to be the most linear), it is still the case that Pokémon are perceived significantly more linear than faces (t(49)=6.9, p<0.0001). Thus both image-based and perceived rectilinearity predict Pokémon selectivity medial to face selectivity.

Next, we generated predictions based on perceived visual features (Methods), using independent raters. In order to get an estimate of how perceived features may affect an untrained visual system, which would mimic the state of experienced participants when they first began playing the game, we chose raters who were not heavily experienced with Pokémon. The first was estimation of ‘real world’ size, which is different from retinal size. For example, a corridor viewed from far away is perceived as large despite subtending a small retinal image. While real world size of faces and corridors are readily estimable, Pokémon do not exist in the real world. For Pokémon, size was evaluated in two ways. First, the game gives information on the physical size of each Pokémon. Second, we evaluated participants’ perceived size of Pokémon (Methods). Twenty-eight participants, who did not participate in the main experiment, were asked to rate the size of various Pokémon using a 1-7 scale, with each number corresponding to reference animals of increasing size. Possible sizes ranged from much smaller than faces (e.g., 1cm; ant) to larger than a corridor (e.g., 7m; dinosaur). Raters’ perception of Pokémon size was largely consistent with the game’s provided physical sizes (average game size: 1.2m±0.95m, average rater size: 0.82m±0.28m). Shown in **Fig 4C**, raters perceive the size of Pokémon to be larger than faces (t(176)=26.3, p<10^−10^), but smaller than corridors (t(176)=16.9, p<10^−10^). As real-world size also shows a lateral (small) vs. medial (large) topography, perceived size predicts that voxels preferring Pokémon in VTC should lie in between face-selective regions on the FG and place-selective cortex in the CoS, that is, on the medial FG.

Lastly, we evaluated the perceived animacy of our stimuli. A separate group of 42 independent participants, rated the perceived animacy of stimuli using a scale of 0-5 (0 most inanimate; 5 most animate). Pokémon are perceived as intermediately animate (**Fig 4D**), less animate than faces (t(82)=13.2, p<10^−10^), but more animate than corridors (t(82)=14.5, p<10^−10^). As animate stimuli are localized to lateral VTC, and inanimate stimuli to medial VTC, this metric predicts that the locus of preferential responses for Pokémon lies, like perceived size, between face- and place-selective cortex on the medial FG.

### Location of novel cortical responses in experienced participants supports eccentricity bias theory

To test these predictions, we produced contrast maps for Pokémon versus all other stimuli in each subject. Using cortex-based alignment (CBA), we transformed each subject’s map to the FreeSurfer average cortical space, where we generated a group-average Pokémon-contrast map. For visualization, we project the map onto an individual experienced subject’s cortical surface. Results reveal three main findings. First, in experienced, but not novice participants, we observed preference to Pokémon reliably localized in the OTS. As illustrated in the average experienced participants’ Pokémon contrast map (**Fig 5A**), higher responses to Pokémon versus other stimuli were observed in the OTS and demonstrated two peaks on the posterior and middle portions of the sulcus. Second, we compared Pokémon activations to those of faces by delineating the peaks of face selectivity in the average contrast maps for faces in each group (**Fig 5A**, white outlines).

**Figure 5:**
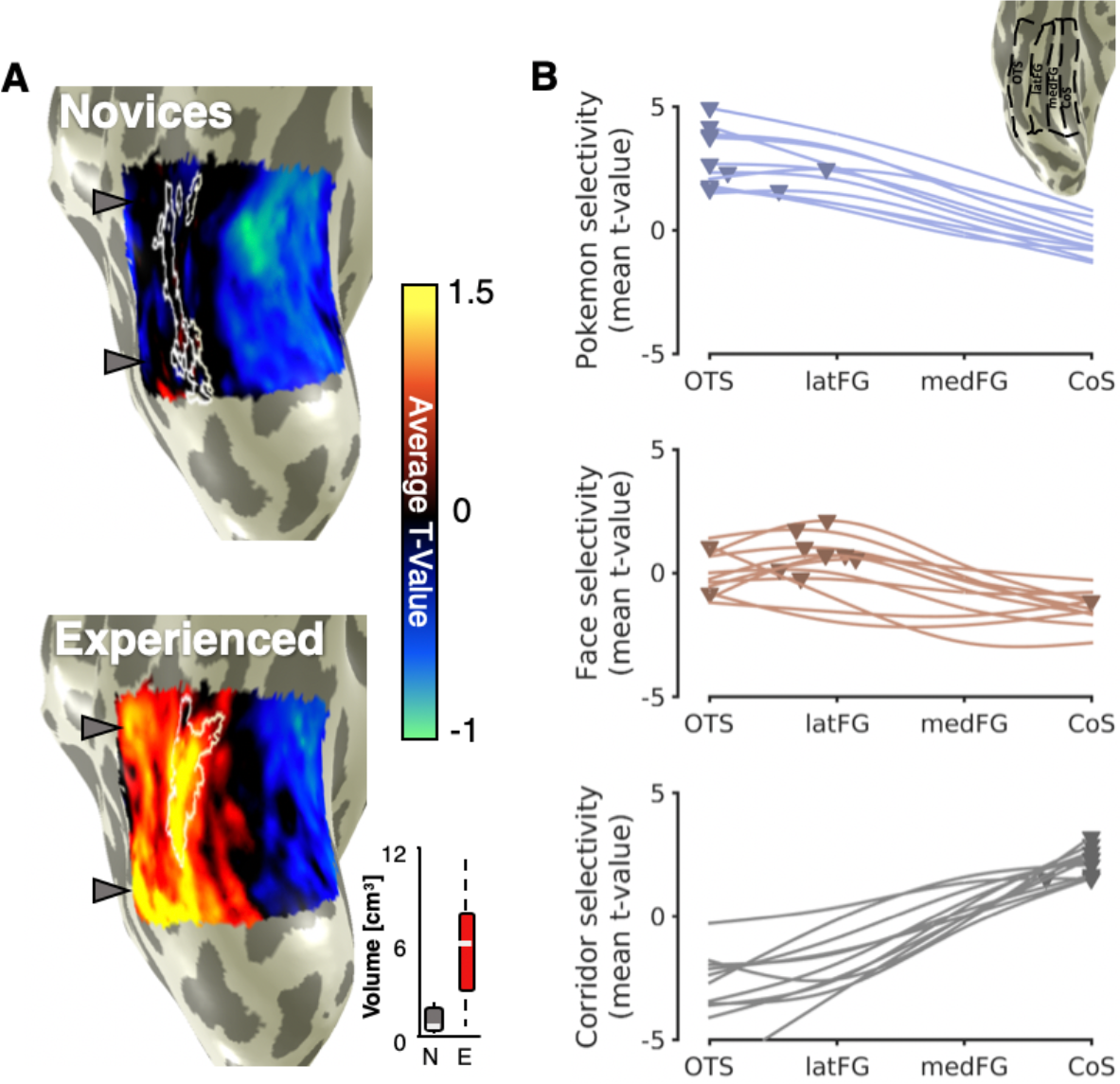
Average contrast maps for Pokémon > all other stimuli in novice and experienced subjects. **A)** In each subject, T-value maps were produced for the contrast of Pokémon versus all other stimuli. These maps were aligned to the FreeSurfer average brain using cortex-based alignment. On this common brain surface we generated a group average contrast map by averaging maps across all novice subjects (left) and all experienced subjects (right). Group average maps are shown on a inflated right hemisphere of one of our participants zoomed on VTC. *White outlines:* group-average face-selective voxels (average T-value > 1) from each respective group. *Gray arrows:* Two peaks in Pokémon-selectivity maps of experienced subjects. The same arrows are shown next to the novice map for comparison. In the lower right, box plots show the mean (white line), 25/ 75% quartiles (boxes) and range (black dotted line) of selectivity volume in novices (N, gray) and experienced subjects (E, red). **B)** Curves fit to the mean selectivity for Pokémon, Faces, or Corridors averaged within one of four anatomically-defined regions extending from lateral to medial VTC (illustrated in the inset for an example subject). Each line is a subject. *Triangle:* peak selectivity value. The peaks of Pokémon-selectivity curves are significantly more lateral than peaks of face-selectivity (t(20)=2.88, p=0.009). The most lateral ROI is the occipitotemporal sulcus (OTS) extending from the ITG to the medial aspect of the OTS. The lateral fusiform gyrus (latFG) ROI includes the lateral FG and ended medially at the mid-fusiform sulcus (MFS). The medial FG (medFG) bin extended from the MFS to the lateral edge of the collateral sulcus (CoS). The collateral sulcus bin (CoS) included the CoS up to the lateral edge of the parahippocampal gyrus.

The comparison reveals that for experienced participants, Pokémon-preferring voxels partially overlapped face-selective voxels on the lateral FG, and extended laterally to the OTS, but never extended medially to the CoS, where place-selective activations occur (**Fig 3**). Third, we compared the volume of category selectivity for Pokémon in each group. Pokémon-selective voxels were any voxels within the VTC that were above the contrast (Pokémon versus all other stimuli) threshold of T-value > 3. Pokémon-selective voxels were observed in the lateral FG and OTS of 11/11 individual experienced participants. In contrast, while some scattered selectivity could be observed for Pokémon in 4 novice participants, it was not anatomically consistent, thus it did not yield any discernible selectivity to Pokémon in the average novice selectivity map (**Fig 5A**). An ANOVA run with factors of group and hemisphere on the volume of Pokémon-selectivity in VTC revealed a main effect of group (F(1,40)=32.75, *p*<0.00001), but no effects of hemisphere nor an interaction (Fs(1,40)<0.67, *ps*>0.41). The median volume in experienced participants was 6-fold larger than novices (**Fig 5A**), with most (7/11) novice participants having near zero voxels selective for Pokémon. This difference in volume between groups was not driven by gender differences (**Fig S2A**). Fourth, we compared the lateral-medial location of Pokémon selectivity relative to face and place selectivity to directly assess theoretical predictions. Thus, we partitioned the VTC in each experienced subject into four anatomical bins from lateral (OTS) to medial (CoS) (**Fig 5B**- inset). Within each bin we extracted the mean t-value for the contrast of either Pokémon, faces, or corridors. Curves fit to average t-values across bins demonstrates that(i) peaks in these curves are the most lateral for Pokémon, intermediately lateral for faces, and medial for corridors (**Fig 5B**), and (ii) Pokémon selectivity peaks are located significantly more lateral in VTC than those of faces (t(20)=2.88, p=0.009). Together, this pattern of results is only consistent with the predictions of the eccentricity bias theory of the development of VTC topography (**Fig 4A**).

Another prediction of the eccentricity bias hypothesis is that the region displaying Pokémon selectivity in experienced subjects should have a foveal bias. A foveal bias predicts that pRF centers will cluster closer to, and result in greater coverage of, the center of the visual field compared to the periphery. This is in contrast to the tiling of the visual field as observed in V1^31,55^, where pRF centers spread across the visual field. To test this hypothesis, six experienced subjects participated in a retinotopic mapping experiment (Methods). Using pRF modeling^56^, we identified for each voxel the region of the visual field which drives its responses. We then generated visual field coverage maps showing how the set of pRFs within Pokémon-selective cortex in each subject tile the visual field. Experienced subjects demonstrate the typical lateral-to-medial gradient of foveal to peripheral bias in VTC (**Fig S5**). In all experienced participants, we find that Pokémon-selective cortex shows a more prominent coverage of central portions of the visual field compared to the periphery. For example, in the right hemisphere of experienced participants, Pokémon-selective cortex shows a contralateral coverage of the visual field, whereby the peak density of the visual field coverage is within 4° of the center of the visual field. With the exception of one subject, the region with the highest pRF density includes the fovea (**Fig 6**). Compared to the eccentricity bias of other categories like faces and corridors, overall Pokémon-selective voxels tend to be the most foveally-biased, followed by face-selective voxels, and lastly corridor-selective voxels, which are the most peripherally-biased (**Fig S6**). Furthermore, the foveal bias of Pokémon-selective pRFs provides evidence consistent with the idea that the eccentricity of retinal images produced by Pokémon biases their cortical responses to emerge in lateral VTC where foveal pRFs exist.

**Figure 6:**
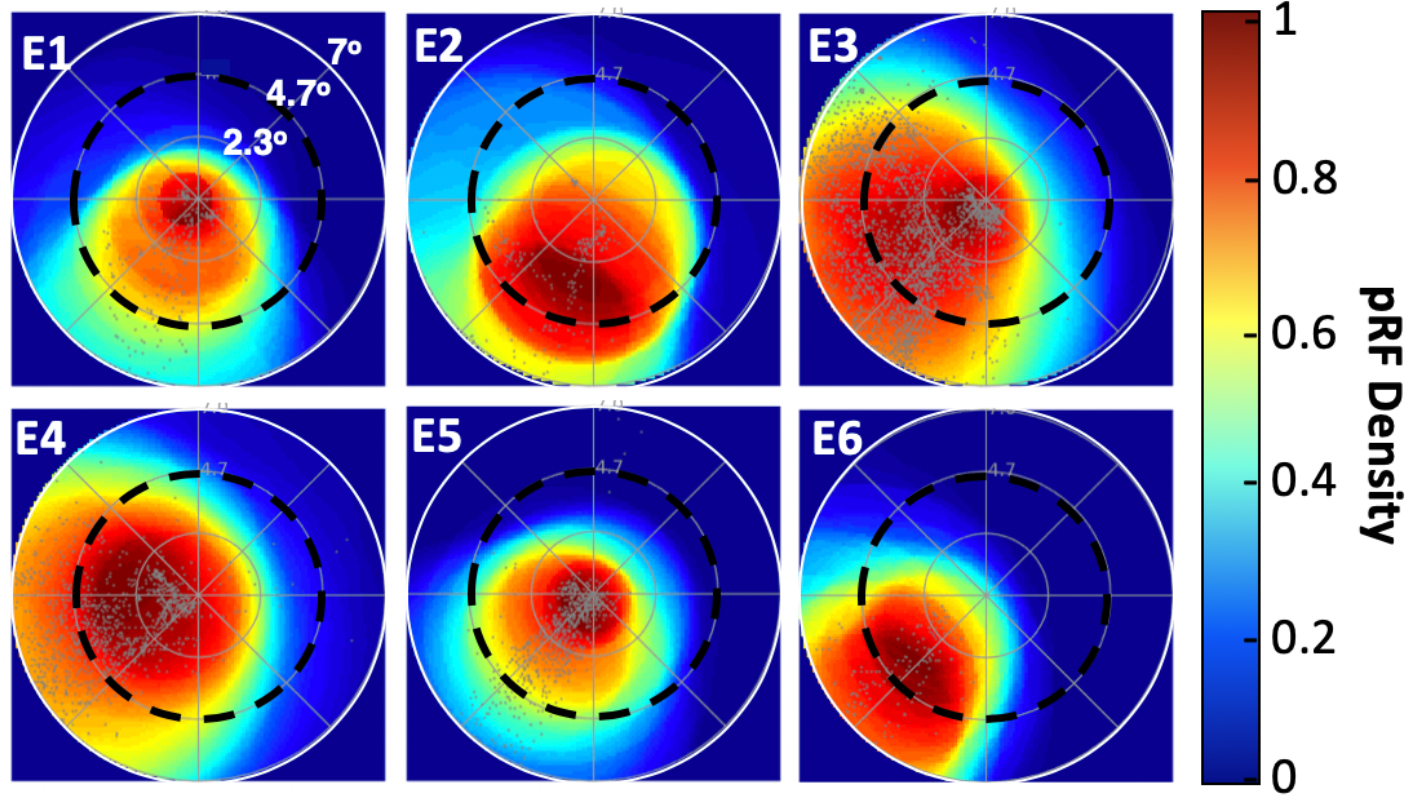
Population receptive field (pRF) modeling reveals Pokémon-selective cortex is foveally-biased. Density plots (see colorbar) representing the visual field coverage by pRFs of Pokémon-selective voxels from the right hemisphere in six experienced subjects. Each panel shows data from a single subject. Density is normalized by the maximum in each subject. *Gray dots:* PRF centers. *Black dashed circle:* 4.7 degrees of eccentricity. Only voxels with greater than 10% variance explained by the PRF model were included.

### How does experience affect the amplitude of responses to Pokémon?

To further understand how novel childhood experience has impacted cortical representations within VTC, we asked two questions: (1) Are the emergent responses to Pokémon in experienced participants specific to the Pokémon characters that participants have learned to individuate, or will similar patterns emerge to any Pokémon-related stimulus from the game? (2) How does visual experience change the responsiveness of OTS to visual stimuli?

To address the first question, a subset of experienced participants (n=5) participated in an additional fMRI experiment in which they viewed other images from the Pokémon game (Methods). In this experiment, participants completed a blocked localizer with two categories of images: images of places (e.g. navigable locations) from the Pokémon game as well as downsampled face stimuli (to resemble 8-bit game imagery). Images were presented in 4 s blocks at a rate of 2 Hz while participants performed an oddball task. Results show that places from the Pokémon game drive responses in the CoS, not OTS or lateral FG. In other words, they produce the typical pattern of place-selective activations (**Fig 7**). In contrast, Pokémon-selective voxels in each subject (**Fig 7**, black contours) have minimal to no selectivity to places from the Pokémon game, further demonstrating the specificity of Pokémon-selective voxels to Pokémon characters.

**Figure 7:**
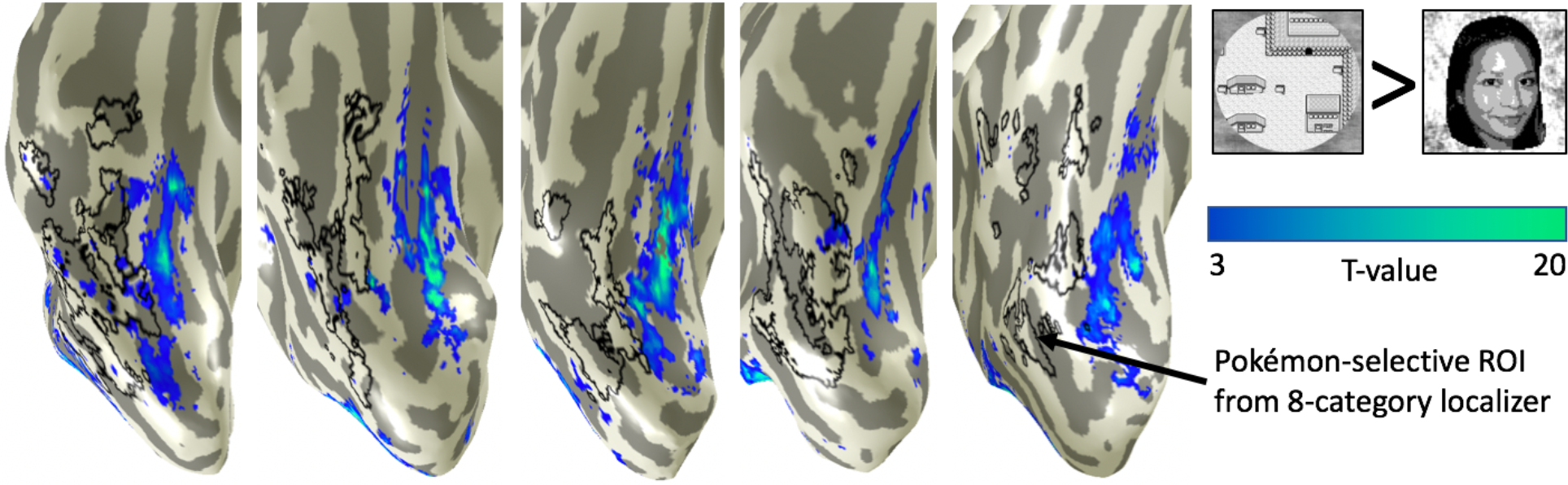
Places from the Pokémon game elicit typical place-selective activations in experienced subjects. Data show T-value maps of Pokémon scenes > pixelated faces in five experienced subjects from a follow-up experiment (see colorbar). A subset of experienced subjects (n=5) participated in an additional fMRI experiment. This blocked experiment contained two categories of interest: downsampled face stimuli (to resemble 8-bit game imagery) and scenes from the Pokémon games. Importantly, these scenes were not required to be individuated within the game. *Black outhne:* Pokémon-selective voxels. Pokémon-selective cortex does not preferentially respond to Pokémon scenes vs. faces; instead Pokémon-selective voxels are in the typical location for place-selectivity, namely, the CoS.

To answer the second question, how experience shapes responses in the OTS, we quantified the response amplitude of Pokémon-selective voxels in both experienced and novice subjects. To ensure that the ROI was defined independently from the individual subject’s data, we employed a leave-one-out approach in which we produced a group-defined Pokémon ROI from ten experienced subjects by transforming individuals ROIs to the FreeSurfer cortical average using cortex-based alignment (CBA) and then used CBA to project the group ROI to the left out subject’s brain and examining its responses. This procedure was repeated for each experienced subject. For novice subjects, we transformed the group ROI produced from all experienced subjects into each individual novice’s brain. To ensure we would not extract signals from cortex that was already selective for another category, in each subject we removed from the group Pokémon ROI any voxels that were selective to other categories. From this independently-defined Pokémon ROI, we extracted the percent-signal change from the 8-category fMRI experiment.

As expected, experienced participants have higher responses to Pokémon compared to other categories (**Fig 8A**, **Fig S7**), with weaker responses to cartoons, animals, and even lower responses to faces, bodies and other stimuli. However, in this putative Pokémon-selective region, novice participants show the highest amplitude of response to animals, then to cartoons and words, and lower responses to the other stimuli (**Fig 8A**). While it may be tempting to conclude that Pokémon-selective voxels emerge from voxels with pre-existing preference to animals, average cortical maps of animal selectivity (**Fig S8**) reveal similar animal selectivity both lateral and medial to face-selective cortex in both novice and experienced participants. Furthermore, perceived animacy ratings (**Fig 4D**) suggest Pokémon should have emerged between face- and place-selective cortex, which would correspond to the portion of the animal selectivity that is medial to face-selective cortex. However, contrary to these predictions, Pokémon selectivity is observed overlapping and lateral to face-selective cortex (**Fig 5**). These results suggest that animacy alone is not driving the emergent locus of Pokémon selectivity.

**Figure 8:**
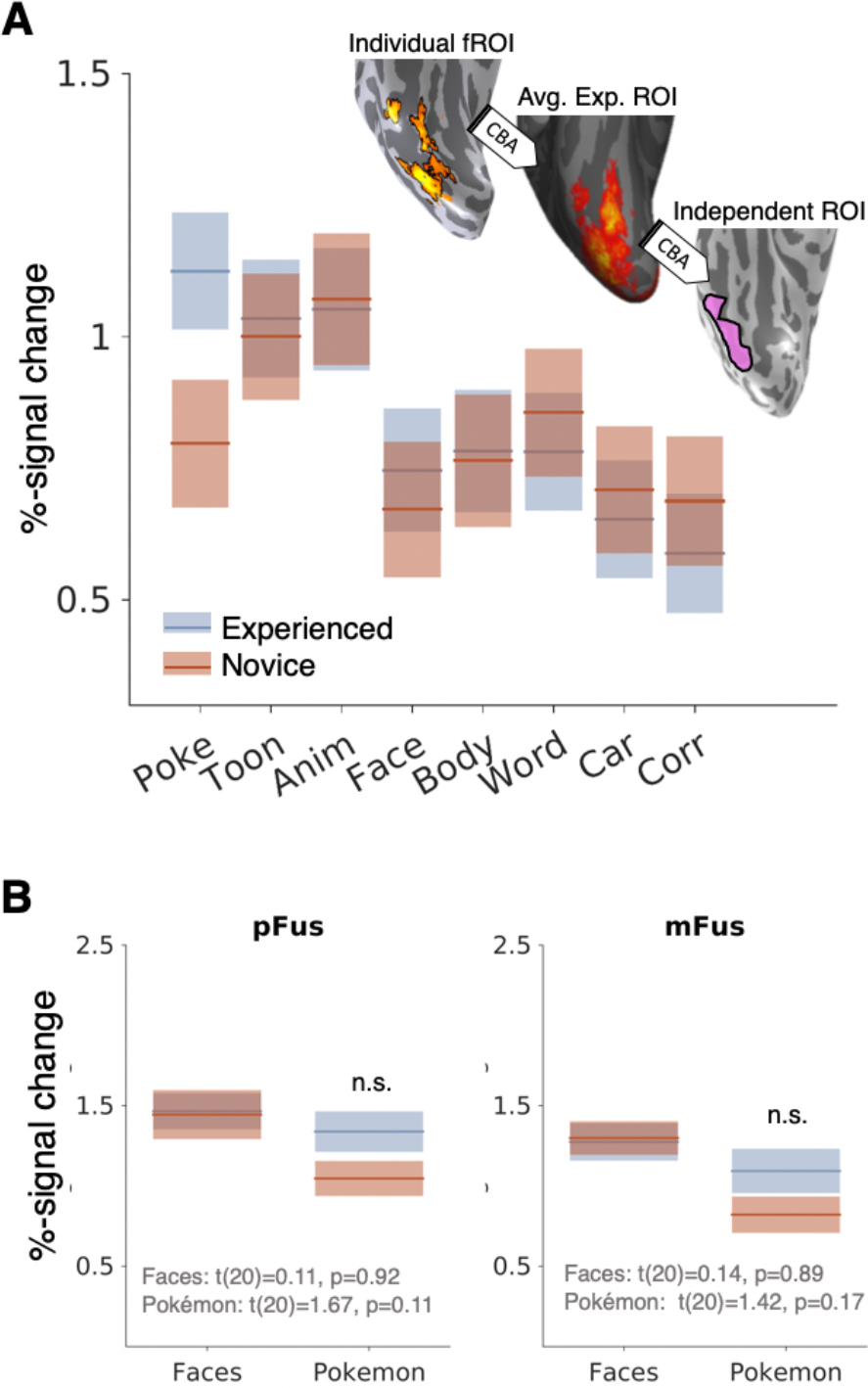
Response properties of the VTC vary with childhood experience with Pokémon. **A)** BOLD responses (%-signal change) measured from an independent definition of Pokémon-selective cortex in both experienced (blue) and novice (orange) participants. **B)** Responses from bilateral pFus- and mFus-faces to faces and Pokémon in experienced (blue) and novice (orange) participants. Face-selective voxels were defined using odd runs of the fMRI experiment, and %-signal change was calculated from the even runs. P-values are from T-tests comparing group differences in responses. Bold line denotes mean, shaded bar is standard error.

To test the expertise hypothesis^49,51^ and because we found that preference to Pokémon stimuli in experienced participants partially overlapped face-selective cortex, we also measured responses to our stimuli in face-selective regions in the posterior FG (pFus-faces) and mid FG (mFus-faces)^57^. We used a split-halves analysis, defining face-selective voxels on the lateral FG from the even runs and extracting the %-signal change from the odd runs to ensure independence of data. Experienced and novice participants did not exhibit differences in the response to faces in either pFus- or mFus-faces, bilaterally (**Fig 8B**). Response amplitudes to Pokémon in pFus- and mFus-faces were numerically higher in experienced compared to novice participants, but not significantly so (**Fig 8B**). Together, analyses of response amplitude illustrate that extensive experience to Pokémon during childhood results in higher responses to Pokémon in adulthood relative to other stimuli leading to the emergence of Pokémon selectivity in and around the OTS and largely outside of other regions of category selectivity.

## Discussion

By examining cortical representations in adults who have undergone lifelong visual experience with a novel visual category beginning in childhood, we offer evidence towards a fundamental question in neuroscience: why is high-level vision organized consistently across individuals? We found that participants who are extensively experienced with Pokémon characters beginning as early as 5 years old demonstrate distinct response patterns in high-level visual cortex that are consistent across participants. Category-selective responses to Pokémon in all experienced participants occupied the posterior and middle extent of the OTS, largely lateral to face-selective cortex, and responded selectively to learned Pokémon characters rather than general imagery from the Pokémon game. Note that we do not assert that our results should be interpreted as a new Pokémon functional module in the OTS of experienced participants on par with face-selective cortex^7^. Rather, our data underscore that prolonged experience starting in childhood can lead to the emergence of a new representation in VTC to a novel category with a surprisingly consistent functional topography across individuals. We demonstrated that this topography is consistent with the predictions of the eccentricity bias theory for two reasons: (i) the small stimuli that required foveal vision during learning biased the emergent representations towards lateral VTC, and (ii) Pokémon-selective voxels in experienced participants show a foveal bias. Together, our data show that shared, patterned visual experience during childhood, combined with the retinotopic representation of the visual system, results in the shared brain organization readily observed in adult high-level visual cortex.

### Experience during childhood generates new brain representations

The nature of Pokémon as a stimulus category is an interesting one, similar to other ecological stimuli such as faces or words, in that the Pokémon game entailed repeated, prolonged, and rewarded experience individuating visually similar, but semantically distinct exemplars. This is similar to other stimulus categories for which there is ecological pressure, or interest, to individuate amongst a visually homogenous category such as faces, birds, or cars^49^. Our data suggest that individuals who have undergone life-long experience individuating Pokémon characters beginning in childhood develop a novel representation for this learned category, demonstrating the plasticity of high-level visual cortex outside of face-selective regions. Specifically, our data show that extensive experience with Pokémon beginning at a young age leads to the formation of a distinct neural representation for Pokémon. This was evinced by robust decoding results and a consistent spatial topography of selectivity for Pokémon in experienced but not novice participants.

Our findings suggest that early childhood visual experience shapes the functional architecture of high-level visual cortex, resulting in a unique representation whose spatial topography is predictable. While one should exercise caution in comparing the current findings, in which participants began their visual experience at a young age, with effects of visual experience in adults^58–60^, differences across studies highlight the possibility that developmental experience may be qualitatively different than that acquired in adulthood. First, childhood experience led to the development of a new representation to Pokémon that was anatomically consistent across participants and was coupled with increased responses and selectivity to Pokémon in the OTS. As the same piece of cortex did not show selectivity in novices, this suggests extensive experience individuating a novel stimulus beginning in childhood is necessary and sufficient for the development of a new representation in VTC. While novices could not name Pokémon like experienced subjects, novices showed similar responses to other unfamiliar categories (e.g. faces, corridors, cars, bodies) that could not be named. Thus, naming ability likely plays little to no role in the group differences we observed. While prior investigations of category training in adults also demonstrated increases in voxel selectivity for the learned category^59,61^, different than our data, these activations were not anatomically consistent across individuals and occurred either in object-selective cortex (LOC) or outside of visual cortex entirely (in the prefrontal cortex). Second, Pokémon elicited numerically but not significantly higher responses in face-selective cortex of our experienced participants. While this data is consistent with prior reports of increased responses in face-selective areas on the FG to stimuli of expertise gained in adulthood^51,58,60^, it is unclear from our data whether these increased responses are due to experience or due to the fact that Pokémon have faces (**Fig S9**). Third, prior research has shown that learning contextual and semantic features of novel objects in adulthood (e.g., this object is found in gardens) can influence VTC representation^62^. Thus, part of the emergent representation for Pokémon in experienced subjects may have stemmed not only from visual features alone, but contextual and semantic information learned about Pokémon. That is, the representation of Pokémon may include additional semantic and contextual information, such as its habitat and characteristics, which can be investigated in future studies. Lastly, developmental work in humans with parametrically morphed stimuli has shown that improved perceptual discrimination among face identities in adulthood is linked with increased neural sensitivity (lesser adaptation) to face identity in face-selective cortex from childhood to adulthood^66^. Higher neural sensitivity is thought to be due to narrower neural tuning. Future research can test if childhood experience with Pokémon also affects neural tuning by measuring adaptation to parametrically morphed Pokémon in experienced versus novice subjects^67^.

Our findings also have interesting parallels with research on the development of reading abilities in children. First, visual experience with Pokémon began between the ages of 5 and 8 years old, similar to ages during which reading ability rapidly improves^63^. Second, it interesting that, like Pokémon, reading words requires foveation and words typically subtend small retinal images. Third, similar to our findings, research on the development of reading has discovered that word-selective cortex emerges during childhood in the lateral OTS and distributed representations in the OTS and lateral FG become more informative from childhood to adulthood with increasing reading experience^64,65^. Thus, our data together with the research on the development of reading, suggest that the critical window for sculpting unique response patterns in human VTC extends to at least school-age.

### Comparison of developmental effects across species

Our results are consistent with previous research in macaques offering compelling evidence for three important developmental aspects of high-level visual cortex: First, its organization is sensitive to the timing of visual experience^43^, as learning new visual categories in juvenile but not adult macaques resulted in new category-selective regions for the trained stimuli. This suggests that there may be critical period during development for cortical plasticity in high-level visual cortex. Second, early visual experience in macaques led to the formation of category selectivity in consistent anatomical locations^33^, and deprivation of visual experience with stimuli like faces results in no face-selective cortex^32^. Third, eccentricity may be a strong prior that constrains development in high-level visual cortex^19,31^. For example, in macaque visual cortex, a proto eccentricity map is evident early in infant development^39,40^.

However, our data also highlight key developmental differences across species. First, the critical window of cortical plasticity in high-level visual cortex may be more extended in humans than macaques. In humans, extensive discrimination training in adults results in changes in amplitudes^58,68^ and distributed representations^59^ in high-level visual cortex, but in adult macaques, responses^43^ and distributed representations^69^ do not change, even as the monkeys become behaviorally proficient at the task, (but see^70^ who showed increases in number of IT neurons responsive to trained stimuli). Second, the anatomical locus of the effects of childhood experience differs across species. In humans, the most prominent functional development have been reported in the FG^20,71–75^ and OTS^65,74^, but in macaques they are largely around the superior temporal sulcus and adjacent gyri^32,33^. Notably, the FG is a hominoid-specific structure^76^, which underscores why development and training effects may vary across species. Third, the features of visual stimuli and how they interact with cognitive strategies that sculpt the brain during childhood may differ across species. That is, developmental predictions about the perceived animacy or size of a visual stimuli is readily queried only in humans.

### Eccentricity as the organizational principle of high-level vision

The unique opportunity presented by Pokémon as a stimulus is the manner in which they vary from other visual stimuli in their physical (retinal image size, rectilinearity) and perceived (animacy, size) properties. Furthermore, the topography of the responses in experienced cortices was consistent across individuals, affording us the ability to ask what potential dimension of Pokémon visual features, either perceived or physical, may determine the anatomical localization of Pokémon responses in VTC. The lateral location of this emergent representation, and its foveally-biased pRFs, suggests the act of foveating on images that subtend a small retinal image during childhood biases input towards regions in lateral ventral temporal cortex which have small pRFs. We posit that individuals experienced with Pokémon had enough patterned visual experience such that this laterally-biased input resulted in category-selectivity.

Several aspects of our data indicate that retinal eccentricity is the dominant factor in determining the functional topography of VTC. While this by no means invalidates observations of other large scale patterns describing the functional topography of high-level vision^5,25,26,35^, our data suggests that retinal eccentricity is a key developmental factor in determining the consistent functional topography of VTC across individuals for several reasons. First, analyses of physical and perceived properties of Pokémon characters suggest that the attribute of Pokémon stimuli that best predicts the location of peak selectivity of responses to Pokémon in VTC was the experienced retinal eccentricity of Pokémon during childhood. Second, pRFs of VTC voxels selective to Pokémon in experienced participants showed a foveal bias. Future research examining how these representations emerge during childhood as participants learn these stimuli will be important for verifying that the foveal bias in the OTS precedes the emerging selectivity to Pokémon. Indeed, while at the image level Pokémon are quite linear in their statistics, individuals are capable of integrating over large pixels holistically to perceive a curve. The extent to which this may be modulated by experience and potentially impact pRFs during learning is an interesting focus of future research. Third, while representation of animacy also exhibits a lateral-medial organization in human VTC, Pokémon are perceived as less animate than faces, but their representation appeared lateral to face-selective regions. In other words, if the animacy axis is continuous across VTC, with animate representations in the OTS and inanimate on the CoS, one would expect Pokémon-selective voxels to be located medial to face-selective cortex. While we observe that such a medial region is capable of responding to animate stimuli (**Fig S8**), it does not become selective for Pokémon across development. Instead, Pokémon-selective voxels were found in the OTS, lateral to face-selective cortex. While high-level visual cortex is capable of distinguishing animate from inanimate stimuli^23^, it might not be a continuous graded representation across the cortical sheet per se. Thus, the framework that human high-level visual cortex has a representation of retinal eccentricity^24,77^, likely inherited from retinotopic input in earlier visual field maps, offers a parsimonious explanation of the development of Pokémon representations in VTC. Future research can examine how becoming a Pokémon expert in childhood, which likely entails learning optimal fixation patterns on Pokémon stimuli, may further sculpt pRFs throughout development as observed in face- and word-selective cortex^31^.

### Conclusions

In a group of individuals who underwent unique, lifelong visual experience beginning in early childhood with a novel visual stimulus—Pokémon—we demonstrate that high-level visual cortex is capable of developing a distinct functional representation for the new learned category. These findings shed light on the plasticity of the human brain and how experience in young age can alter cortical representations.

An intriguing implication of our study is that a common extensive visual experience in childhood leads to a common representation with a consistent functional topography in the brains of adults. This suggests that how we look at an item and the quality with which we see it during childhood affects the way visual representations are shaped in the brain. Our data raise the possibility that if people do not share common visual experience of a stimulus during childhood, from either disease, as is the case in cataracts^28,78^, or cultural differences in viewing patterns^79^, then an atypical or unique representation of that stimulus may result in adulthood, which has important implications for learning disabilities^80,81^ and social disabilities^82^.

In sum, our study underscores the utility of developmental research, showing that visual experience beginning in childhood results in functional brain changes that are qualitatively different from plasticity in adulthood. Future research examining the amount of visual experience necessary to induce distinct cortical specialization and determining the extent of the critical window during which such childhood plasticity is possible will further deepen our understanding of the development of the human visual system and its behavioral ramifications.

## Acknowledgements

We thank 3 anonymous reviewers for their constructive feedback. This research was funded by the Ruth L. Kirschstein National Research Service Award grant F31EY027201 to JG, and NIH grants 1ROI1EY02231801A1 and 2RO1 EY 022318-06 to KGS.

## Author contributions

JG designed conducted, and analyzed the data in this study, as well as wrote the manuscript. MB designed and conducted the study.

KGS oversaw the study, data analyses, and wrote the manuscript.

## METHODS

### Subject details

Human participants were between the ages of 18 and 44, with a mean and standard deviation of 26.8 ± 4.8 years. Participants were split into two groups: experienced (n=11; age 24.3 ± 2.8, 3 females) and novice participants (n=11; age 29.6 ± 5.4, 7 females). The former group was selected initially through self-report, with the inclusion criteria that (1) participants began playing the original Nintendo Pokémon games between the ages of 5 and 8 on the handheld GameBoy device, (2) continued to heavily play the game and its series throughout their childhood, and (3) either continued to play the game into adulthood or revisited playing the game at least once in adulthood. Novice participants were chosen as individuals who never played the Pokémon game and had little to no interaction with Pokémon otherwise. All participants completed a 5-choice naming task designed to test the naming ability of participants; experienced participants scored significantly better than novices (**Fig 1A**). All participants provided written consent to participate in the experiment per Stanford University IRB.

### Behavioral Naming Task

Participants completed a 5-choice naming task with 40 items of randomly selected Pokémon. The experiment was self-paced and participants were told to choose the correct name of each Pokémon. Performance was evaluated as %-correct accuracy, with experienced subject significantly outperforming novices, two-tailed t-test t(18)=17.3,p<10^−11^. The behavioral identification task was run after scanning had been completed in order to minimize exposure of Pokémon stimuli to novice participants. Two novices and one experienced subject were unable to complete behavioral testing.

### fMRI 8 category experiment

All participants underwent functional magnetic resonance imaging (fMRI), completing six runs of the experiment, with different images across runs. Each run was 218 seconds in duration, and across the six runs, counterbalanced 4s blocks presented 8 stimuli at 2Hz. Categories included faces, headless bodies, 8-bit Pokémon sprites from the original Nintendo game, animals, popular cartoon characters from television of the late 1990s and early 2000s, pseudowords, cars, and corridors. Stimuli, as illustrated in **Fig 1B**, were presented on a textured background produced by phase scrambling a stimulus from another category^83^. Stimuli and backgrounds were counterbalanced so that all stimuli appeared with all other phase-scrambled categories as a background. This was done such that the Fourier amplitude spectra of all images across categories will be matched as much as possible, as done in prior publications^31,66,83^. Stimuli were presented approximately in the center of the background square but were jittered in size and position to minimize differences in the average image between categories. Stimuli were run through the SHINE toolbox^84^ to remove luminance differences between categories or potential outlier stimuli. Animals (which includes insects) were chosen as an animate stimulus category that closely resembles Pokémon, as most Pokémon characters were designed to resemble an animal or insect. Cartoons were chosen because they are another animated category that was experienced by both groups predominately in childhood and was somewhat recognizable by both groups.

### fMRI Pokémon scenes and pixelated faces control experiment

To evaluate that the information we observed in the OTS in experienced participants for Pokémon was selective for the learned game characters and not just any stimulus from the game, a subset of experienced participants (n=5, 3 of which also complete pRF mapping detailed in the following section) was invited back to the lab to complete a follow-up fMRI experiment containing two categories of images. The first consisted of scene stimuli from the Pokémon game. These scenes were map-like images extracted from the game through which the player has to navigate their character, including towns, natural landscapes, and indoor spaces. These scenes were free of any images of people or Pokémon. The second category of stimuli were human faces (from the main experiment) except that they were downsampled to visually resemble the 8-bit pixelated images from the Pokémon game in order to match for low-level visual features. This was accomplished by resizing face stimuli to 70×70 pixels (approximately the original pixel size of Pokémon characters) and then resized again using bicubic sampling to their original size of 768×768 pixels. Images were presented in 4s blocks at a rate of 2hz and were interleaved with a blank baseline condition. There were 12 blocks of each condition in a run; each run was 162s seconds in durations and participants completed 3 runs. Data were analyzed similarly to the main category localizer. Participants were instructed to pressing a button when a blank image appeared within a block of images (oddball detection task).

### Population receptive field (pRF) mapping

To evaluate the visual field coverage of receptive fields within Pokémon-selective cortex, 5 experienced participants from the main experiment were invited to complete a pRF mapping experiment. Each participant completed 4 runs of a sweeping bar stimulus that traversed the 7×7 degrees-of-visual-angle screen in eight different directions, as implemented in previous work^31^. For each voxel, the pRF model^56^ was fit using a circular gaussian with a position (x,y), size (sigma), and a compressive spatial summation^85^ to account for BOLD nonlinearities beyond V1. For each subject, Pokémon-selective cortex was defined as any voxel in lateral VTC demonstrating a T-value of 3 or greater for the contrast of Pokémon versus all other stimuli from the localizer data. To derive pRF density maps to understand how the visual field is covered by the pRFs of each subject’s Pokémon-selective ROI (**Fig 6**), pRFs were plotted as circles using the fit size (standard deviation of the fit pRF gaussian) and the visual field was colored pointwise according to pRF density, where 1 corresponds to the max pRF overlap in that subject.

### Behavioral 2-back task

To ensure that novice participants are capable of visually detecting differences between Pokémon characters, and thus demonstrate that they don’t perceive all Pokémon as an indistinguishable homogenous object, we invited a separate group of novice participants (n=36) to the lab in order to complete a 2-back repetition detection task. Subjects saw blocked images from one of three categories (faces, Pokémon, corridors) and were instructed to press a button when they detected an image repeat with an intervening image. Stimuli were presented at 2Hz and 8 images were presented within a block, as was done during functional MRI. There could be 0 or 1 repeats within a given block, and 50% of blocks contained a repeat; there were 40 blocks per category. Blocks with repeats were randomly assigned and were counterbalanced across categories, and category and stimulus order were randomized.

### Anatomical MRI

T1-weighted anatomical volumes of the entire brain were collected for each subject at a resolution of 1mm isotropic at the end of each functional scanning session. Anatomical scans were acquired with a T1-weighted BRAVO pulse sequence with the following parametersinversion time = 450ms, flip angle of 12°, field of view = 240mm. Anatomical volumes were processed using FreeSurfer^86^, https://surfer.nmr.mgh.harvard.edu. Volumes were segmented to gray and white matter to produce a reconstruction of the cortical surface as well as a definition the cortical ribbon (gray matter). Functional data were restricted to the cortical ribbon.

### fMRI acquisition

T2*-weighted data were collected on a 3 Tesla GE Discovery scanner using a gradient echo simultaneous multi-slice (SMS) acquisition protocol for 3x simultaneous slice readout with CAIPI-z phase shifting to improve SNR of the reconstructed image^87^. Slices (16 prescribed, 48 in total after SMS) were aligned parallel to the parieto-occipital sulcus (POS) in each subject to guarantee coverage of the occipital and ventral temporal lobes. Parameters were as followsFOV = 192mm, TR= 2, TE = 0.03s, voxel size 2.4mm isotropic, no interslice gap.

### fMRI data processing

Functional data were motion-corrected within and between scans and aligned to an anatomical reference volume per the standard Vistasoft pipeline^75^, see https://github.com/vistalab/vistasoft. Localizer data were unsmoothed and always analyzed in native subject space (unless producing average cortical maps as described below). Functional data were corrected for within and between-scan motion; all participants were well-trained and moved less than a voxel through their scans. For each voxel, the timecourse was transformed from arbitrary scanner units to percent-signal change (PSC) by dividing every timepoint by the mean response across an experiment. Localizer data were fit with a standard general linear model by convolving stimulus presentation times with a difference-of-gaussians hemodynamic response function implemented in SPM (www.fil.ion.ucl.ac.uk/spm). The GLM were also used to perform contrast maps between different stimulus conditions.

### Multivoxel pattern analyses (MVPA)

We performed MVPA on ventral temporal cortex (VTC) voxel data. VTC was defined in each subject’s cortical surface using the following cortical folds as in our prior publicationslaterally, it was bounded by the occipitotemporal sulcus (OTS) up to the beginning of the inferior temporal gyrus (ITG), medially, by the collateral sulcus (CoS), anteriorly, by the anterior tip of the mid-Fusiform sulcus (MFS), and posteriorly, by the posterior transverse collateral sulcus (ptCoS). The multivoxel patterns (MVPs) in response to each category (betas resulting from the GLM) were represented as a vector, the values of which were z-scored by subtracting the voxel’s mean response and dividing by 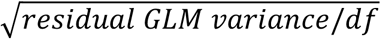. We then computed all pairwise correlations between MVPs of one category to every other to produce the 8×8 representational similarity matrices in **Fig 2A**. Each cell is the average correlation from split-half combinations of the 6 runs (e.g. 1-2-3/4-5-6, 2-3-4x/1-5-6…). MVPAs were run on in-plane data using the original fMRI data in the acquired resolution.

### Classification

To quantify the information present within MVPs of VTC, we constructed a winner-take-all (WTA) classifier. The classifier was trained from one half of the data and tested how well it could predict the held-out half of data. For a given split half we used one half as a training set and the other half as the testing set. The WTA computes the correlation between the MVP of given test data (MVP to an unknown stimulus) and each of the MVPs of the labeled training data. It classifies the test data based on the training data that yielded the highest correlation with the test data. For a given test MVP, correct classification yields a score of 1, and an incorrect choice yields a score of 0. Performance was computed for each category for both permutations of training and testing sets, yielding possible scores of 1 (both classifications were correct), 0.5 (if only one was correct), or 0 (both incorrect). For each category, we averaged classification scores from all split-half combinations (ten) per participants, and then averaged classification performance across participants within a group to produce the values in **Fig 2B,C.**

### Measurement of mean ROI response amplitudes

To evaluate the BOLD response from Pokémon-selective cortex in experienced participants independently from the data used to define the ROI, we used FreeSurfer to produce an average Pokémon-selective ROI that was independent of each subject’s individual data. We implemented a leave-one-out approach, defining for the n-th subject an average Pokémon ROI from n-1 subjects. In 10 experienced subjects, we transformed voxels selective for Pokémon (Pokémon vs. all other stimuli, t>3, voxel level) using cortex-based alignment to the FreeSurfer average cortical surface (an independent average of 39 individuals). We then generated a group average probability map of the location of Pokémon on the FreeSurfer average brain and thresholded maps from these 10 experienced participants to only include vertices that were consistent across at least 30% of participants. This threshold has been employed in previous research and demonstrates the optimal point in Dice coefficients for predicting ROI location in human cortices^75,88^. Furthermore, the thresholding ensures that no one subject in particular can influence the group average ROI. This thresholded group ROI was then transformed back into the left-out participant, from which we then extracted the %-signal change reported in **Fig 8A**. This procedure was repeated 11 times, for each of the experienced subjects. In a similar manner, we created on the FreeSurfer average brain a group Pokémon ROI from all 11 experienced subjects thresholded at the same 30% overlap level. This group ROI was transformed using CBA into the each of the novice participants’ brains, from which we extracted the %-signal change to our stimuli. Because CBA is not as accurate as defining fROIs from data in each subject’s brain, we excluded voxels that showed significantly selectivity to another category

### Definition of pFus- and mFus-faces

Face-selective cortex was defined in each subject’s brain as in prior work using cortical folds within VTC as anatomical landmarks. mFus-faces was a cluster of face-selective (T-values greater than 3 for the contrast of faces versus all other stimuli) voxels located on the lateral FG, aligned to the anterior end of the mid fusiform suclus (MFS). pFus-faces was defined with the same contrast and was located on the lateral FG 1-1.5cm posterior to mFus-faces.

### Evaluating the anatomical peak location of face, Pokémon, and corridor selectivity in VTC

The goal of this analysis was to determine the location of peak responses to a given category on a linear (lateral to medial) space to test the theoretical predictions illustrated in Fig 4. Thus, in each experienced participant, we defined four anatomical bins in VTC arranged from lateral to medial, whose posterior and anterior extent are constrained by the posterior transverse collateral sulcus and the anterior tip of the mid-Fusiform sulcus (MFS), respectively. The lateral and medial edges of each bin are as followsi) the lateral-most bin extended from the inferior temporal gyrus (ITG) through the occipitotemporal sulcus (OTS) and ended at the fusiform gyrus (FG). ii) The next bin included the lateral FG and ended at the pit of the MFS. iii) The next bin extended from the MFS and included the medial FG, ending at the collateral sulcus (CoS). iv) The most medial bin included the CoS and ended at the parahippocampal gyrus. Example bins are shown in the inset of **Fig 5B**. From each bin, and within each subject separately, we extracted the mean selectivity (t-values) for a given contrast (faces, Pokémon, or corridors). For each category we then fit spline curves across these four values to produce the curves shown in **Fig 5B**. Spline curves were fit in order to allow for smoother spatial information and also aided in more accurately identifying peak locations. From these curves, we identified the maxima in each subject, and compared the lateral-to-medial coordinate values across categories.

### Image statistics analyses

To generate predictions for the locus of Pokémon-selective voxels in VTC compared to face- and place-selective voxels, we ran a number of analyses on images of Pokémon, faces, and corridors. We evaluated two physical properties of the stimulieccentricity of the retinal image, and curvilinearity of the images, and two perceived propertiesphysical size and animacy of the stimuli. Corridors were chosen as they represent a category of scene stimuli that all participants are likely to have equal visual experience, and because they are matched to Pokémon stimuli in that they possess linear features. This provides a larger dynamic range between faces and scenes to more readily assess where Pokémon exist in visual feature space between these stimuli classes.

#### Eccentricity bias

To evaluate the average eccentricity of the retinal image, we simulated retinal images (144 per category) in a representation of the visual field spanning 150 by 150 degrees of visual angle (75 degrees in one hemifield) and used a distribution of likely viewing distances for stimuli to encapsulate the variability of viewing distance in real life and individual differences in viewing behavior. For corridors and rooms, we assumed the image nearly always occupied the entire visual field. That is, corridor/room stimuli were simulated to occupy anywhere from 95 to 100% of the visual field by assigning a random value 95 to 100% the size of the 150° simulated visual field (i.e. 75° radius) to produce a distribution for the retinal image size of places. For stimuli of faces and Pokémon, we simulated Gaussian distributions of retinal image sizes. For Pokémon, viewing distance is somewhat fixed, as they are viewed on the GameBoy screen (characters are presented in a 2cm square, and on average occupy 1.6cm on the GameBoy screen) and limited to a maximum of arm’s length. We assumed a relaxed position in which a subject would, on average, hold the device 32cm^90^ from the eyes with a standard deviation of 10cm. Pokémon were also viewed in childhood in the cartoon show, which aired on TV once a week for a duration of 12-14 minutes. Thus, we estimate much less time was spent watching Pokémon on TV than playing the GameBoy game. Bernard Lechner of RCA laboratories measured average viewing distance at which individuals watched television (national average of 9 feet), and produced optimal viewing distances for television screens of a given resolution. Given Lechner’s distance for average TV viewing, and the size of TVs in the late 1990s and early 2000s, the retinal image produced by Pokémon from the cartoon show is comparable to the closely held GameBoy. For faces, which have an average size of 20cm, and an average viewing distance of 150cm (about 5 to 6 feet)^91^, we used a standard deviation of 50cm in viewing distance. While individuals may fixate on a face at further (or very close distances), these viewing times are likely far less in magnitude than the conversational viewing distance of a face which likely composes the majority of face-viewing time. The average size of a face was taken to be 20cm. Each stimulus of each category was then simulated as a retinal image from which we calculated its eccentricity in visual degrees, assuming that participants fixated on the relevant item. randomly sampled from the distributions described above. The distribution of these eccentricity values are displayed in **Fig 4A**.

#### Rectilinearity

The lines composing a given image can be quantified in terms of their rectilinearity, that is, how straight (linear) or rounded (high curvature) are the composite features of an image. We employed the rectilinearity toolbox^25^, which convolves a series of wavelet filters varying in orientation (22.5°–360° in 22.5° steps), position, angle (30°, 60°, 90°, 120°, 150°, and 180°), and scale (1/5, 1/9, 1/15, and 1/27 cycles per pixel) to assess the rectilinearity of our images. All Pokémon, face, and corridors stimuli were analyzed by this toolbox, whereby each image is given a score it relative to other stimuli. The output of this toolbox is a ranking of all provided stimuli from least to most linear. Histograms based on the distribution of stimulus categories in this linear ranking are plotted in **Fig 4B**.

### Perceived attributes

#### Size

Faces and places have distinct and measurable sizes in the environment, and producing distributions around the mean size of a face or place is relatively straightforward. For places, we will be using corridors in our simulations as their size, compared to something open air like a forest or beach, has a readily estimable size. Given that open air places are likely larger than the numbers simulated here, these simulations thus represent a conservative estimate for the size of place stimuli. For faces and places, we simulate distributions of their real-world sizes, and later for Pokémon, which have no real-world size, we will have raters report their perceived size and compare these values to those of faces and places. Assuming an average anatomical head size of 20cm in height, we make a Gaussian distribution with a standard deviation of 2cm around that point. Distribution sample size was 150. The average wall height in a standard storied building is 8 feet, but can be up to 9 or 10 feet. Corridors and building interiors likely make up a large percentage of place interactions for our participants, however, we also account for some larger scenarios that might be encountered in larger buildings or sky scrapers. We thus made a skewed distribution with a tail towards larger sizes; no value for corridor size was allowed to go below 2.43m. Distribution sample size was 150. For Pokémon, which are modeled within the game to have animal-like properties, have explicit sizes within the game itself. For example, a given Pokémon is accompanied by information about its physical features, including its size. While viewers experience the Pokémon on a small screen (similar to viewing images of faces and corridors on a small screen), there is an inherent understanding of what its physical size would be in the real world. To evaluate the physical size of Pokémon we took two approaches. First, we used the actual in-game sizes as the distribution of physical size. Second, we had an independent group of participants (n=28) rate what they thought the size of (n=50) Pokémon randomly selected from the 144 stimuli were using a 1-7 Likert scale. Each integer corresponded to a reference animal of increasing size (ant, mouse, cat, dog, gorilla, horse, dinosaur). Raters’ choices were converted from a likert digit (2, corresponding to a mouse), to the average metric size of that animal (mouse = 7.5cm). We found that independent raters perceived Pokémon to be of sizes strikingly similar to those within the game. Each rating (Likert integer) was converted into meters using the average size of the reference animal, and a mean size was calculated for each subject. Participants’ ratings were then compared to the distribution of face sizes (n=150) and corridor sizes (n=150) from above.

#### Animacy

To quantify animacy for our images, we had another independent group of raters (n=42) evaluate the perceived animacy of Pokémon, faces, and corridors on a scale of 1 to 5, with increasing values corresponding to higher animacy. Participants were shown one image at a time, and directed, for each image, the following: “How animate, or living, do you perceive this image to be?” Participants rated 50 stimuli each of faces, Pokémon, and corridors; stimuli from each category were randomly selected from the total 144 stimuli used in the localizer experiment for each category. For each subject, their mean rating of animacy was calculated by averaging their scores for faces, Pokémon or words. Distributions were then made over the raters’ mean values for each stimulus category.

#### Rectilinearity

To quantify how individuals may perceive different stimuli as more or less rectilinear, we conducted another behavioral experiment in an independent group of raters (n=50). In the experiment they viewed images of faces, Pokémon characters, and corridors, and evaluated on a scale from 1 (very curvy) to 7 (very linear/boxy) how linear they perceived the stimuli to be. The experiment was self-paced, and participants were instructed to examine the lines, edges, and shapes that made up a particular image and evaluate overall how linear they perceived the particular object to be. Participants saw 40 images of each category. From each subject we derived a mean rating for each category; Violin plots depicting the distribution across subjects for the ratings of faces, Pokémon, and corridors are plotted in **Fig S4**.

## Data Availability

Further information and requests for resources should be directed to and will be fulfilled by the Lead Contact, Jesse Gomez

## Supplementary Figures

**Supplementary Figure 1:**
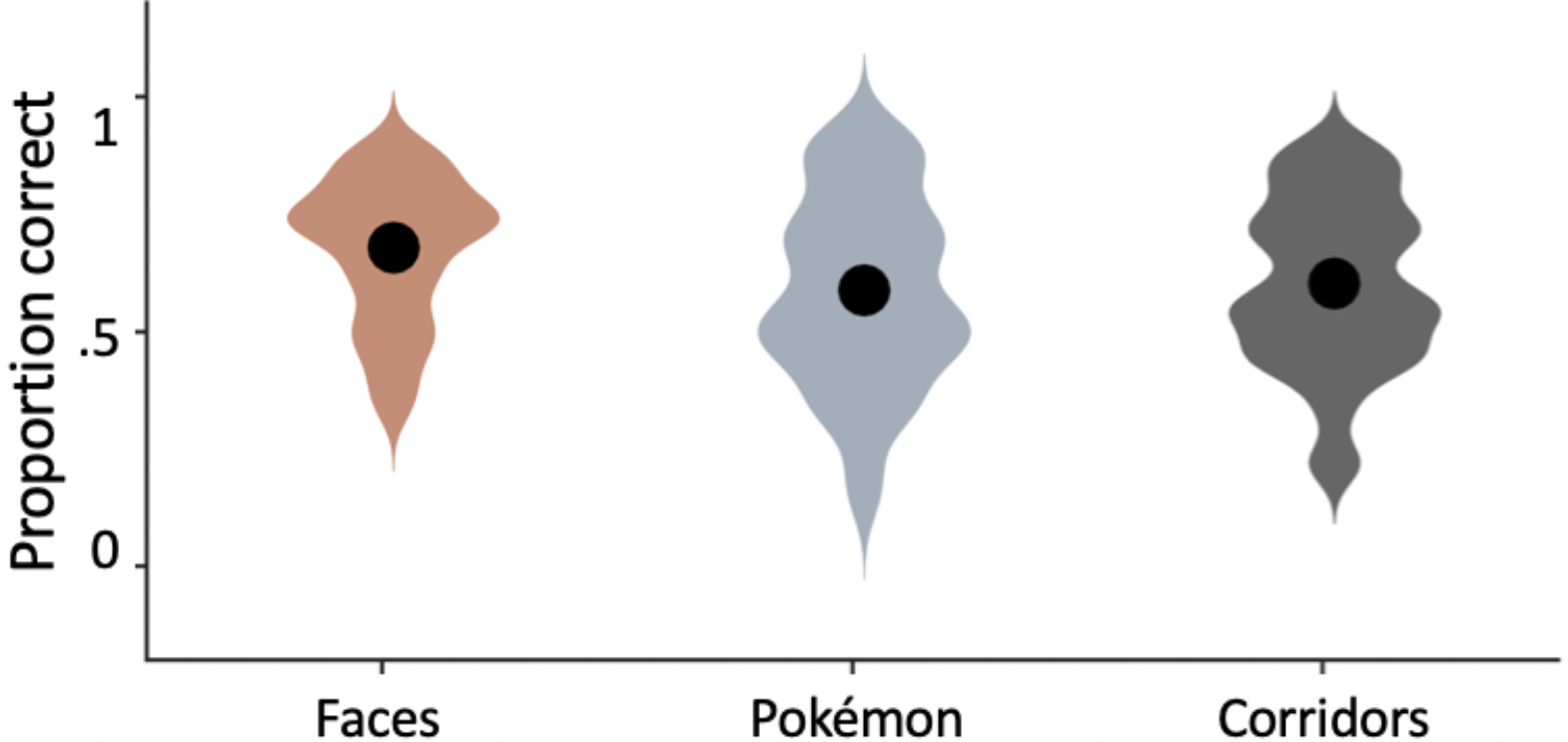
Detection accuracies for the 2-back task by stimulus category. Participants’ accuracies for detecting a 2-back repeat during the 2Hz presentation experiment. Violin plots depict the distribution of n=36 individuals in their accuracy scores for detecting repeats of faces (orange), Pokémon (blue), and corridors (gray). Large black circles illustrate the mean rating across all participants. Novice participants were similarly accurate on the 2-back task on Pokémon as on corridors (t(34)=0.5, p=0.62), but lower than faces (t(34) = 3.3, p=0.002).

**Supplemental Figure 2:**
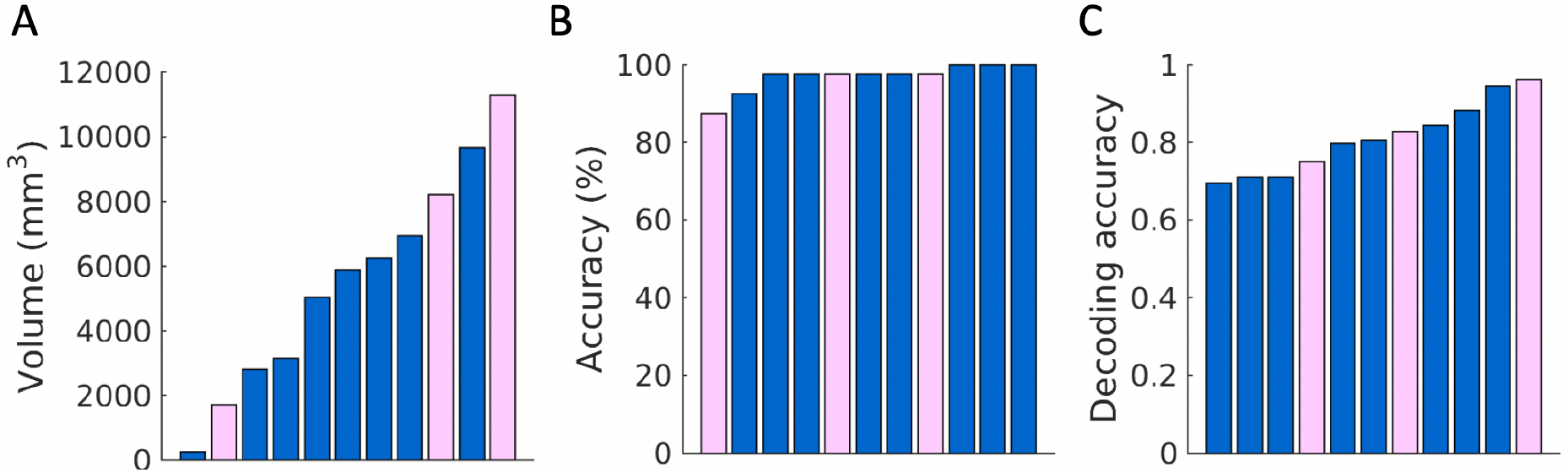
Participants experienced with Pokémon who differ in gender do not differ in effect sizes. (A) Bar plots depicting the volume of t>3 voxels for the contrast of Pokémon versus all other stimuli VTC for all experienced subjects. Females, shown in pink, are distributed evenly across subjects and are not biased towards one extreme. (B) Bar plots depicting the %-correct accuracy on the Pokémon-naming behavioral experiment in experienced subjects. Females (pink) are evenly distributed. (C) Bar plots of the decoding accuracy in experienced subjects of the winner-take-all classifier show in Figure 2 by individual subject. Again, females are evenly distrbuted amongst all subjects.

**Supplemental Figure 3:**
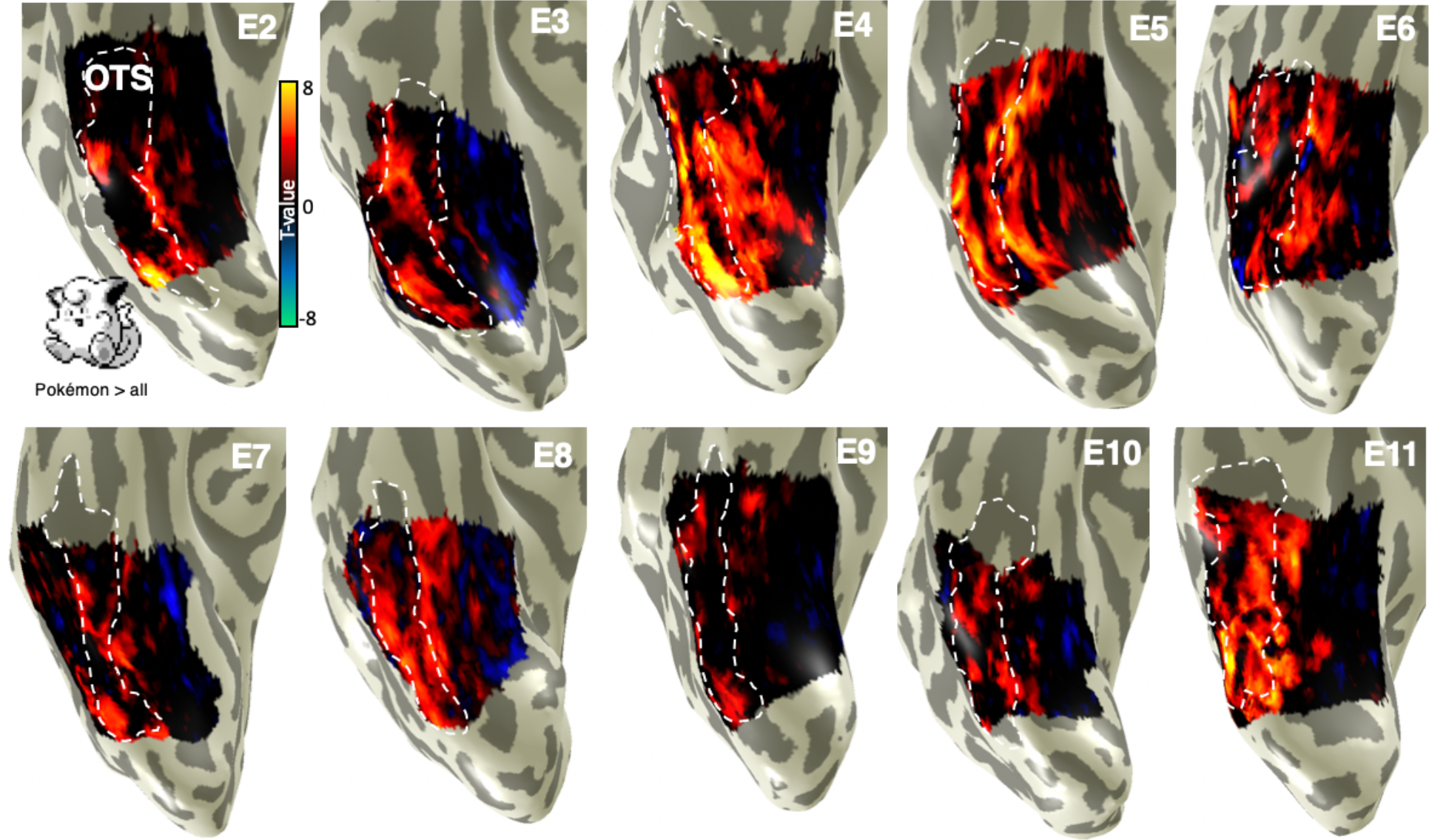
Distinct cortical representation for pokémon in experienced subjects. Unthresholded parameter maps for the contrast of Pokémon versus all Other stimuli in all subjects. Each panel shows the inflated ventral right cortical surface zoomed on VTC. *White dotted line:* OTS All subjects demonstrate Pokémon-selectivity Within the posterior arri middle OTS.

**Supplementary Figure 4:**
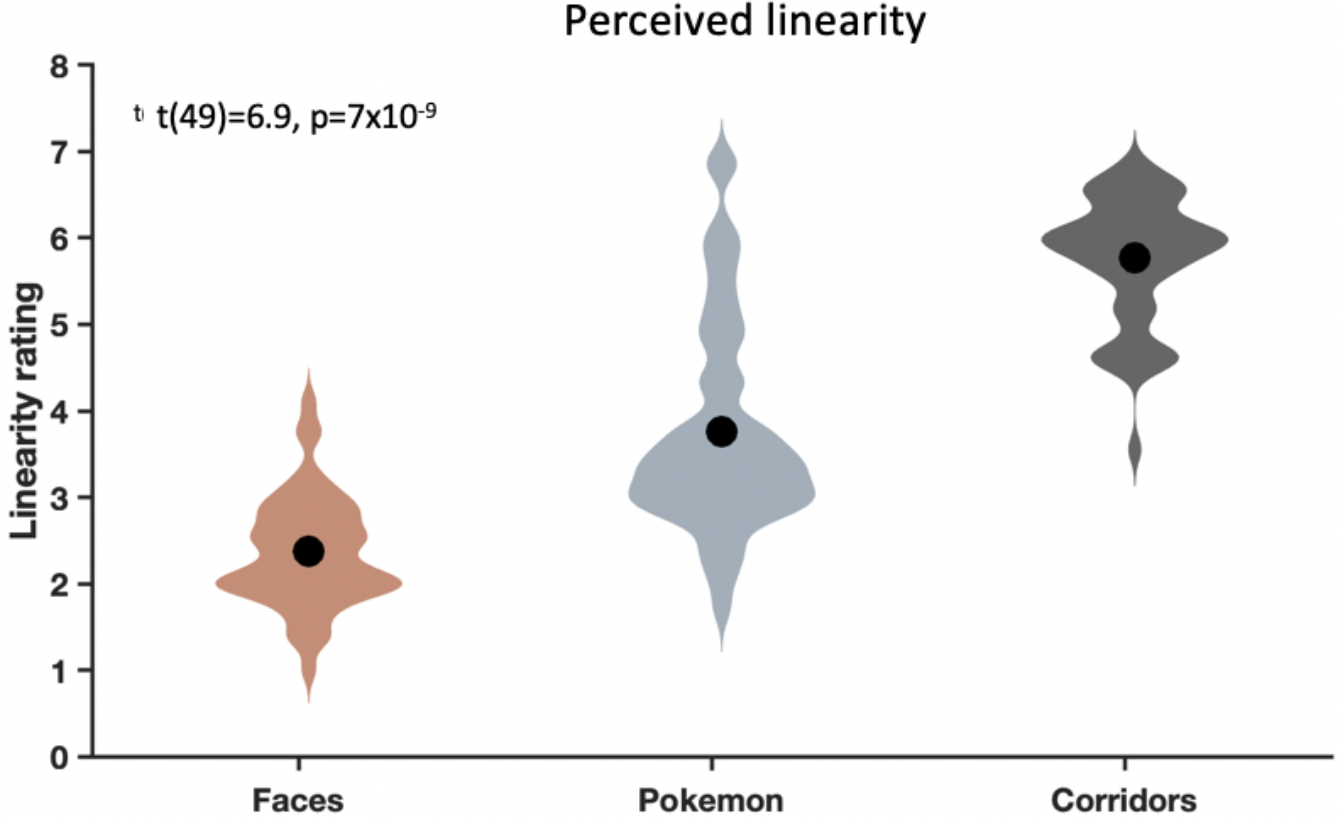
Ratings of perceived linearity for faces, Pokémon, and corridors. N=50 subjects completed a behavioral rating experiment in which they were asked to rate how curvy/round (score of 0) to linear/boxy (score of 7) a given stimulus appeared to be. For each participant, an average value was calculated for faces, Pokémon, and corridors. Violin plots depict subject density across the 50 participants in their mean ratings for a given stimulus category. Large black circles illustrate the mean rating across all participants. T-statistic represents a paired t-test comparing the linearity scores given be subjects to faces versus Pokémon. Pokémon are rated a significantly more linear than faces.

**Supplementary Figure 5:**
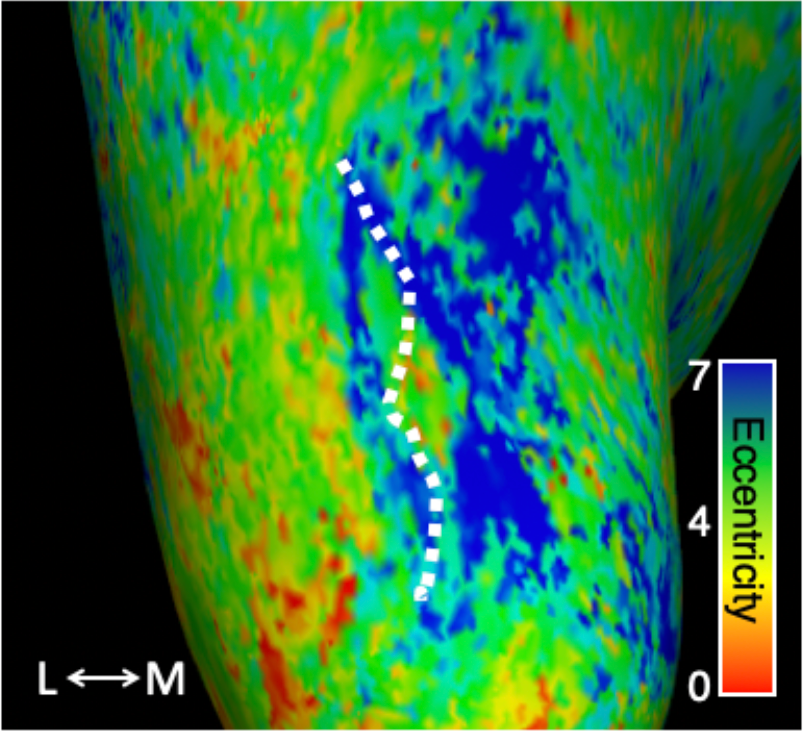
Experienced subjects show typical eccentricity gradient in VTC. Group-average eccentricity map from six experienced subjects who underwent pRF mapping. Experienced subjects demonstrate the typical eccentricity gradient in VTC, with more foveal representations lateral to the MFS (dotted-white outline) and more peripheral representations medial to it. Colorbar shows average pRF eccentricity in degrees of visual angle. *L:* lateral. *M:* Medial.

**Supplementary Figure 6:**
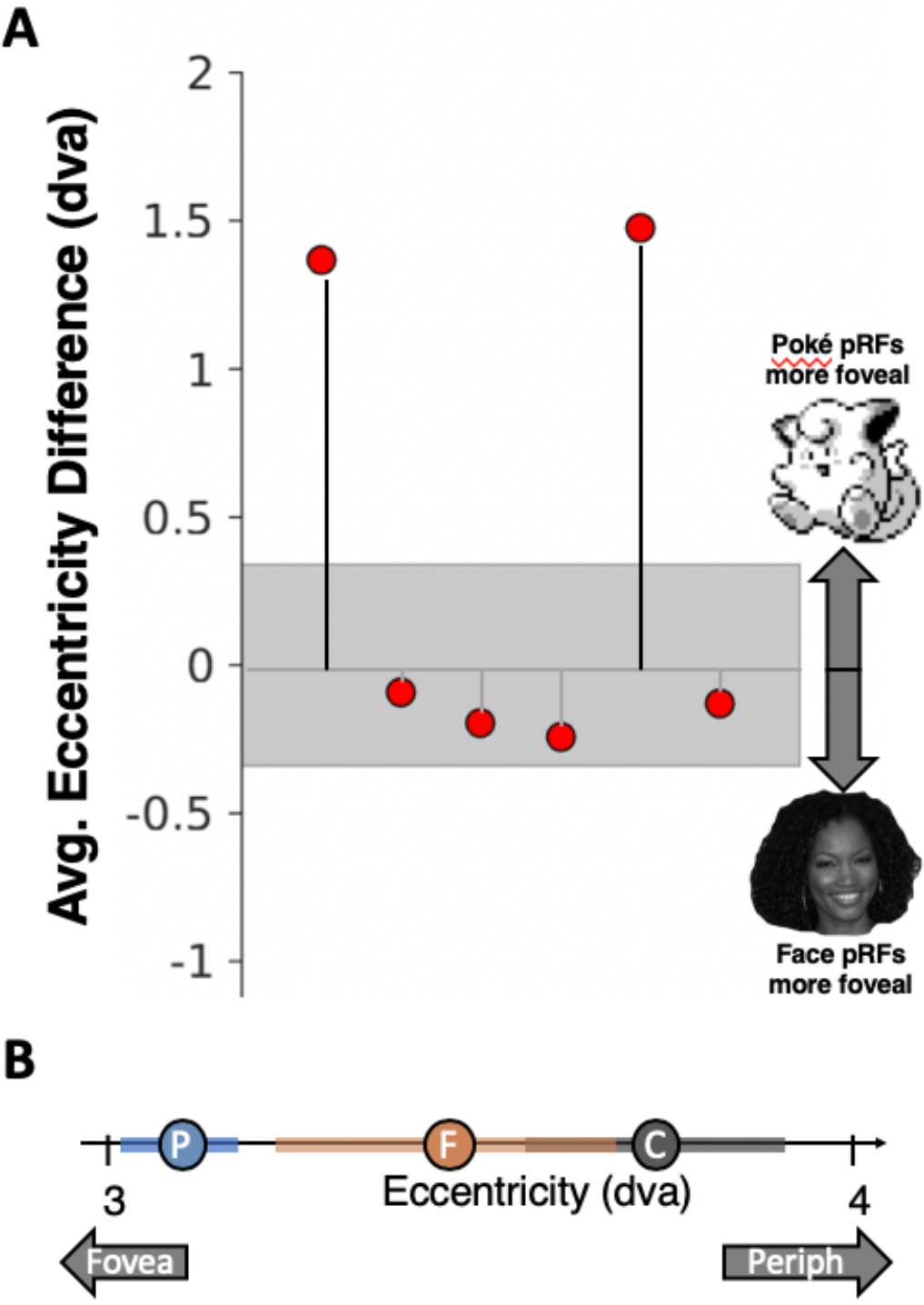
Relative foveality bias between face-selective and Pokémon selective voxels in the VTC of six experienced subjects. (A) For each of 6 subjects, the average eccentricity of voxels that were either face-selective (t>3) or Pokémon-selective (t>3) was calculated. Then a difference score in each subject was derived, such that positive values indicate (in degrees of visual angle, dva) how much closer to the center of the visual field Pokémon-selective pRFs were relative to face-selective pRFs. Gray region depicts standard error across subjects. (B) Circles and standard error bars depicting the mean distance of pRF centers across subjects for Pokémon-selective voxels (blue), face-selective voxels (orange), and corridor-selective voxels (gray). Circles are plotted on a single line depicting distance from the fovea in degrees of visual angle (dva).

**Supplemental Figure 7:**
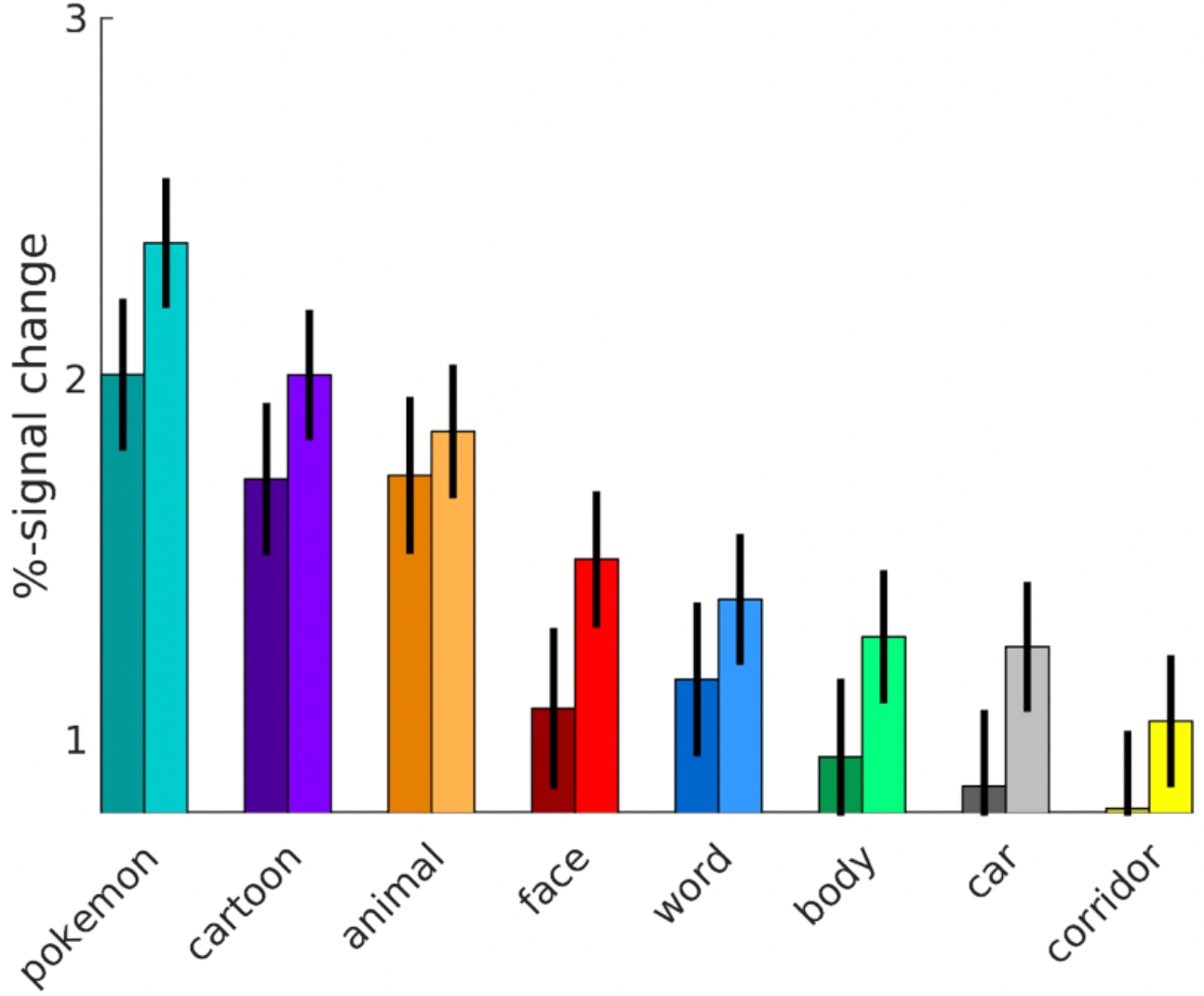
Responses from each experienced subject’s Pokémon-selective ROI in the OTS. This is non-independent data and only meant to illustrate the responses from each subject’s ROI that contribute to category selectivity. *Dark bars:* left hemisphere; *Light bars:* right hemisphere. Despite being in the OTS where the visual word form area can be found, Pokémon-selective voxels’ response to words is low, on par with other inanimate categories such as cars and corridors.

**Supplemental Figure 8:**
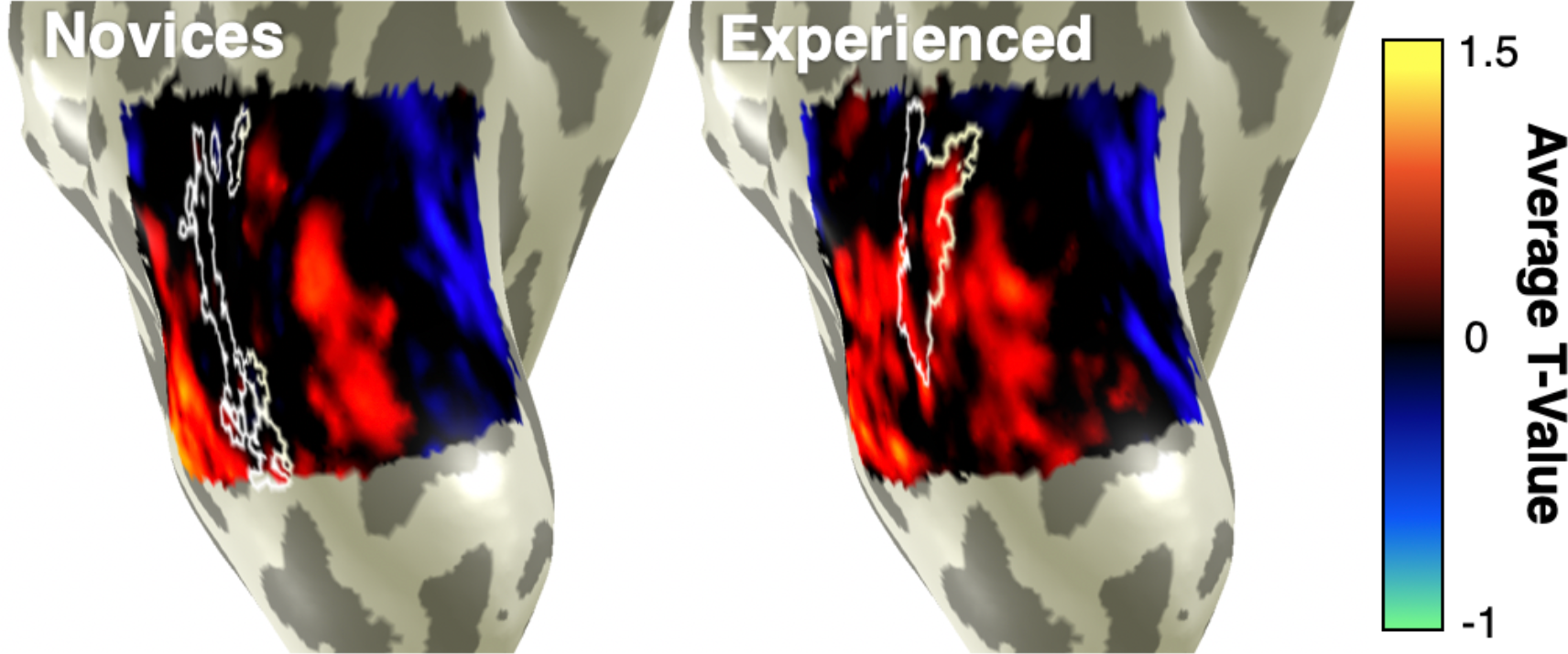
Group-average contrast for animals versus all other stimuli in novice and experienced subjects. Average contrast maps for the preferred category of animals were produced in each individual subject and then averaged using cortex-based alignment in FreeSurfer. Average face-selective cortex in each group is outlined in white. In both subject groups, animal-selectivity can be seen both medial and lateral to face-selective cortex. Data are shown on an example inflated right hemisphere zoomed on VTC. Same subject as Fig. 5.

**Supplementary Figure 9:**
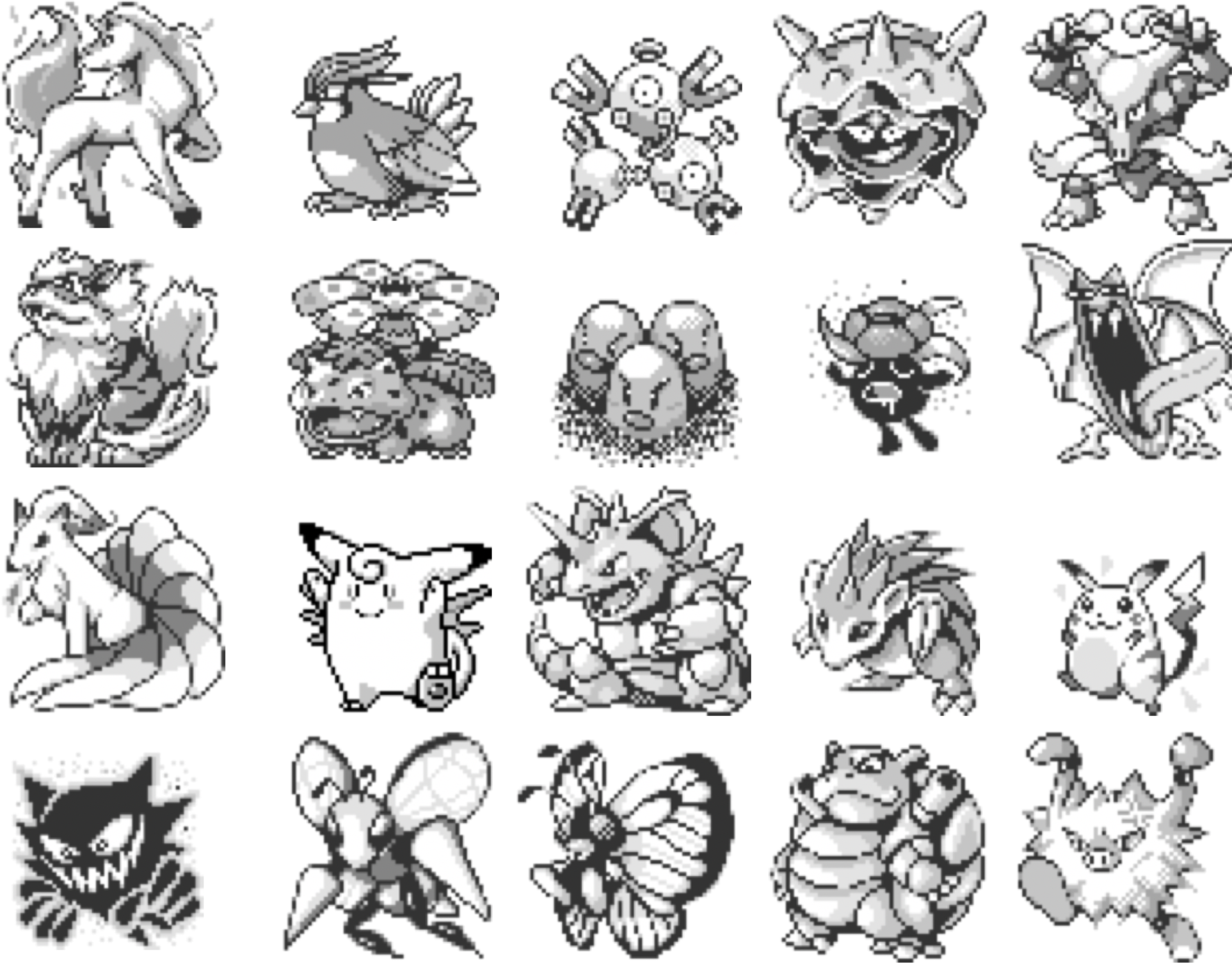
Example Pokémon Characters. Pokémon were designed, in most cases. to resemble animals. and for the must part have readily faces features such as feet, arms, legs, hands.

